# Visibly Transparent, Near-Infrared Absorbing Nanofluids Enable High-Efficiency and Safe Laser Lithotripsy

**DOI:** 10.64898/2026.05.31.729132

**Authors:** Qingsong Fan, Junqin Chen, Arpit Mishra, Megan Bock, Aaron Stewart, Catherine Cai, Judith Dominguez, Jiadong Liu, Yuanke Chen, Ronghui Wu, Ting-Hsuan Chen, Jiaoti Huang, Christine Payne, Micheal E. Lipkin, Pei Zhong, Po-Chun Hsu

## Abstract

Laser lithotripsy (LL) is the gold standard for urinary stone management yet maximizing ablation efficiency while maintaining procedural safety remains clinically challenging. Here, we present a visibly transparent, near-infrared (NIR)-absorbing ITO@SiO_2_ nanofluid irrigation strategy that significantly enhances LL efficiency without compromising endoscopic visibility. By spectrally matching the absorption profile of ITO@SiO_2_ with the clinical Holmium:YAG laser wavelength, ablation efficiency improved by >200% in the bench-top spot treatments and >100% in the hydrogel kidney model. Mechanistic investigations revealed that the enhanced optical absorption of the nanofluid modifies bubble dynamics and synergistically amplifies photothermal/microexplosion effects and cavitation damage. Importantly, both in-vitro hydrogel and in-vivo porcine kidney models demonstrated a substantial thermal safety margin (maximum temperatures <35 °C) and excellent acute biocompatibility, with no evidence of thermal tissue injury. Integrating seamlessly into established clinical workflows without requiring stone pretreatment, this strategy offers a highly translatable, safe, and efficient platform for next-generation endoscopic lithotripsy.

## Introduction

Urinary stone disease is a globally prevalent urological disorder (**Fig. 1a**), with incidence peaking between the third and fifth decades of life.^1–3^ Multiple studies have highlighted its increasing prevalence in recent decades, driven largely by metabolic and lifestyle-related risk factors including dietary changes, obesity, and type 2 diabetes.^2^ In the United States (U.S.), the lifetime prevalence of stone disease increased from 3.2% in 1980 to 10.1% in 2014.^4^ Furthermore, retrospective analyses of U.S. commercial and Medicare populations from 2011 to 2019 revealed that diagnosed cases increased from 1.84 million to 2.09 million, translating to an annual incidence rate exceeding 1.3% by 2019 (**Fig. 1b**).^5^ This growing disease also imposes substantial economic costs on healthcare systems, with annual expenditure for urolithiasis in the U.S. projected to reach $4.6 billion by 2030.^6^ Consequently, according to the Urologic Diseases in America project, urolithiasis now stands as the costliest urologic condition in the nation, surpassing even the aggregate costs of other major diseases such as prostate cancer.^5^

**Fig. 1.**
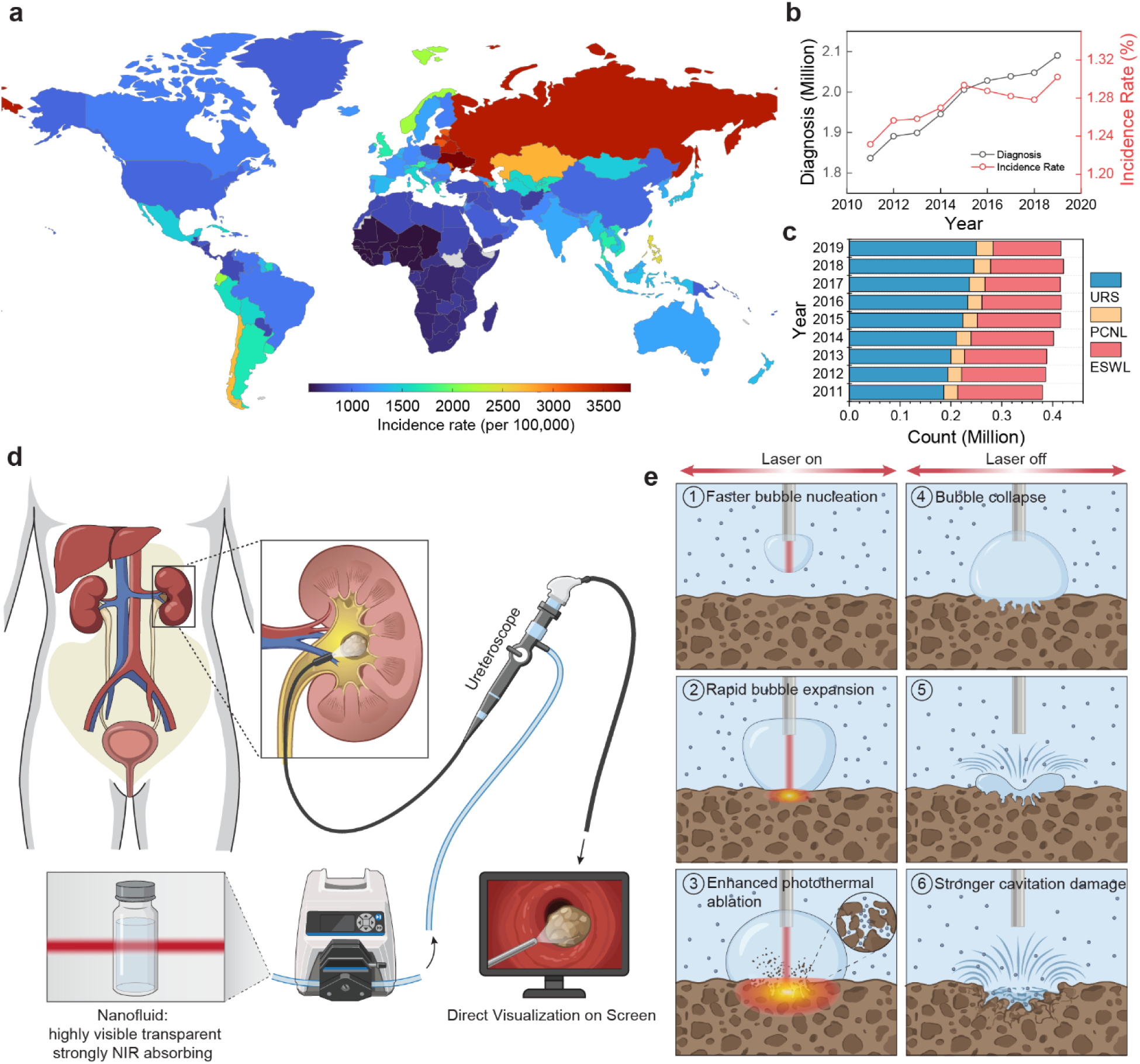
(a) Global incidence of acute urolithiasis in 2021 (Data adapted from Ref.^3^) (b) Annual diagnoses and incidence rates of urinary stone disease in the U.S. from 2011 to 2019 (Data adapted from Ref.^5^). (c) Annual procedural volumes of ESWL, URS, and PCNL in the U.S. from 2011 to 2019. (d) Schematic illustrating the integration of a visibly transparent, near-infrared (NIR)-absorbing ITO@SiO_2_ nanofluid into standard laser lithotripsy (LL) procedures. (e) Proposed mechanism of nanofluid-enhanced stone ablation involving accelerated bubble dynamics, enhanced photothermal/microexplosion effects, and intensified cavitation damage.

Patients experiencing an acute episode of urinary stone disease often suffer from debilitating pain, severe nausea, and renal dysfunction.^7^ Renal colic caused by kidney stones is widely regarded as one of the most severe pains reported, with some educational sources cite a pain score of 42 out of 50 for kidney stones, above childbirth and tooth pain.^8,9^ Fortunately, with the advancement of technology, patients now have multiple treatment options. The most widely recommended interventions include extracorporeal shock wave lithotripsy (ESWL), ureteroscopy (URS), and percutaneous nephrolithotomy (PCNL). ESWL is a non-invasive procedure that utilizes high-energy shock waves to fragment stones. Conversely, URS is a minimally invasive technique in which a thin, flexible ureteroscope is passed through the urethra, bladder and distal ureter, enabling the urologist to see and treat stones *in situ* in the upper urinary tract under direct vision. For larger or more complex stones, PCNL involves inserting a nephroscope through a small percutaneous incision in the back to break up and extract the stone. **Supplementary Table 1** and **2** compare these three modalities across clinical metrics such as recurrence rate, complication rate, contraindications, and average hospital cost.^10^ Notably, URS has gained increasing clinical preferences owing to its minimal invasive nature, broad applicability, shorter recovery times, lower recurrence rates, and limited contraindications, making it a uniquely viable option for pregnant patients and those with bleeding diatheses.^11^ Consequently, URS procedural volumes have steadily increased in the U.S., while the use of ESWL has declined (**Fig. 1c**).^5^ This shift toward URS is part of a broader global trend, observed in countries including the United Kingdom, New Zealand, Australia, Canada, and Brazil.^12^

Historically, a variety of lithotripters have been employed for stone fragmentation, including ultrasonic, electrohydraulic, and pneumatic devices, as well as various laser systems (e.g., pulsed ruby, dye, and Q-switched Nd:YAG lasers) during URS.^1,13^ However, due to the significant limitations of these earlier modalities (**Supplementary Table 3**), lithotripters other than the Holmium:YAG (Ho:YAG) laser are rarely used today. Owing to its versatility and reliable performance across stones of varying compositions, the Ho:YAG laser has become the clinical gold standard for ureteroscopic lithotripsy.^14^

Operating at a wavelength of 2120 nm, the Ho:YAG laser is strongly absorbed by water, enabling it to outperform other lasers operating in the 500 – 1000 nm and effectively ablate stones of all compositions via two primary mechanisms:^13,15^ (1) Photothermal effect: when the laser energy reaches the stone through the Moses effect, a portion of the light is absorbed and converted into heat, leading to localized melting, thermal decomposition, and weakening of the stone matrix. In addition, water confined within the micropores of the stone can undergo rapid vaporization, generating internal microexplosions that further promote stone disintegration, and (2) Photoacoustic effect: recent studies have demonstrated that ring-like cavitation damage can be produced by the shock wave emission from the toroidal bubble collapse around the laser ablation spot. Recently, the Thulium Fiber Laser (TFL) has emerged as a promising alternative.

Operating at 1940 nm, the TFL targets a wavelength where water’s absorption coefficient is roughly four times higher than that of the Ho:YAG laser.^16^ Although numerous benchtop experiments have reported superior dusting efficiency using TFL, robust in-vivo validation and comprehensive safety assessments remain limited.^17–19^ Consequently, the Ho:YAG laser continues to represent the clinical standard for ureteroscopic lithotripsy.

Enhancing the efficiency of the Ho:YAG laser not only shortens operative times but also reduces the costs associated with equipment upgrades (such as transitioning to the TFL), thereby mitigating the overall financial burden on healthcare systems. To date, the majority of research efforts have focused on manipulating the laser’s output profile—such as pulse energy, pulse duration, pulse modulation, and frequency.^20,21^ In contrast, comparatively little attention has been directed toward engineering the optical absorption environment surrounding the laser–stone interaction zone, despite its central role in governing both cavitation dynamics and photothermal energy transfer.

Several previous studies have explored the use of absorptive nanoparticles including gold, carbon, or hollow Prussian blue nanoparticles, to modify the stone surface and enhance laser absorption.^22,23^ However, these approaches remain poorly suited for clinical translation. This is primarily due to a mismatch between the nanoparticles’ absorption peaks and the wavelength of the Ho:YAG laser (**Supplementary Table 4**). Moreover, many reported strategies require direct pretreatment or coating of the stone surface, which is incompatible with routine endoscopic workflows.

To address this gap, herein we propose a novel workflow that is highly compatible with current clinical practices for Ho:YAG laser lithotripsy (LL). Specifically, we developed a nanofluid composed of ITO@SiO_2_ nanoparticles that exhibits strong absorption near Ho: YAG wavelength while maintaining minimal attenuation in the visible range (**Fig. 1d**). Unlike previous methods that require pretreatment of the stone surface, our approach directly replaces standard saline irrigation with this engineered nanofluid during the LL procedure. Using this strategy, we demonstrate substantial enhancement of stone ablation efficiency through synergistic amplification of photothermal ablation/microexplosive effects and cavitation-mediated damage (**Fig. 1e**).

## Results

As an n-type semiconductor, ITO nanoparticles feature free charge carriers that enable localized surface plasmon resonance (LSPR) under incident electromagnetic fields. In contrast to noble metals (e.g., Au and Ag) that absorb strongly in the UV-visible region, the relatively lower carrier concentration in ITO shifts its LSPR peak into the near- or mid-infrared range (**Fig. 2a**). This unique profile perfectly matches the Ho:YAG laser wavelength. Furthermore, modifying the Sn doping level directly alters the carrier concentration, enabling tuning of the corresponding LSPR peak position. As shown in **Fig. 2b**, undoped In_2_O_3_ nanoparticles exhibit weak overall absorption with no distinct peak within the measured wavelength range. However, upon tin doping, a strong absorption peak emerges, which gradually shifts from ∼2300 nm to ∼1800 nm as the feeding ratio increases from 3.23% to 7.69%. Notably, this absorption peak is strictly confined to the NIR region, ensuring that the solution remains highly transparent even at high concentrations due to its negligible absorption in the visible spectrum.

**Fig. 2.**
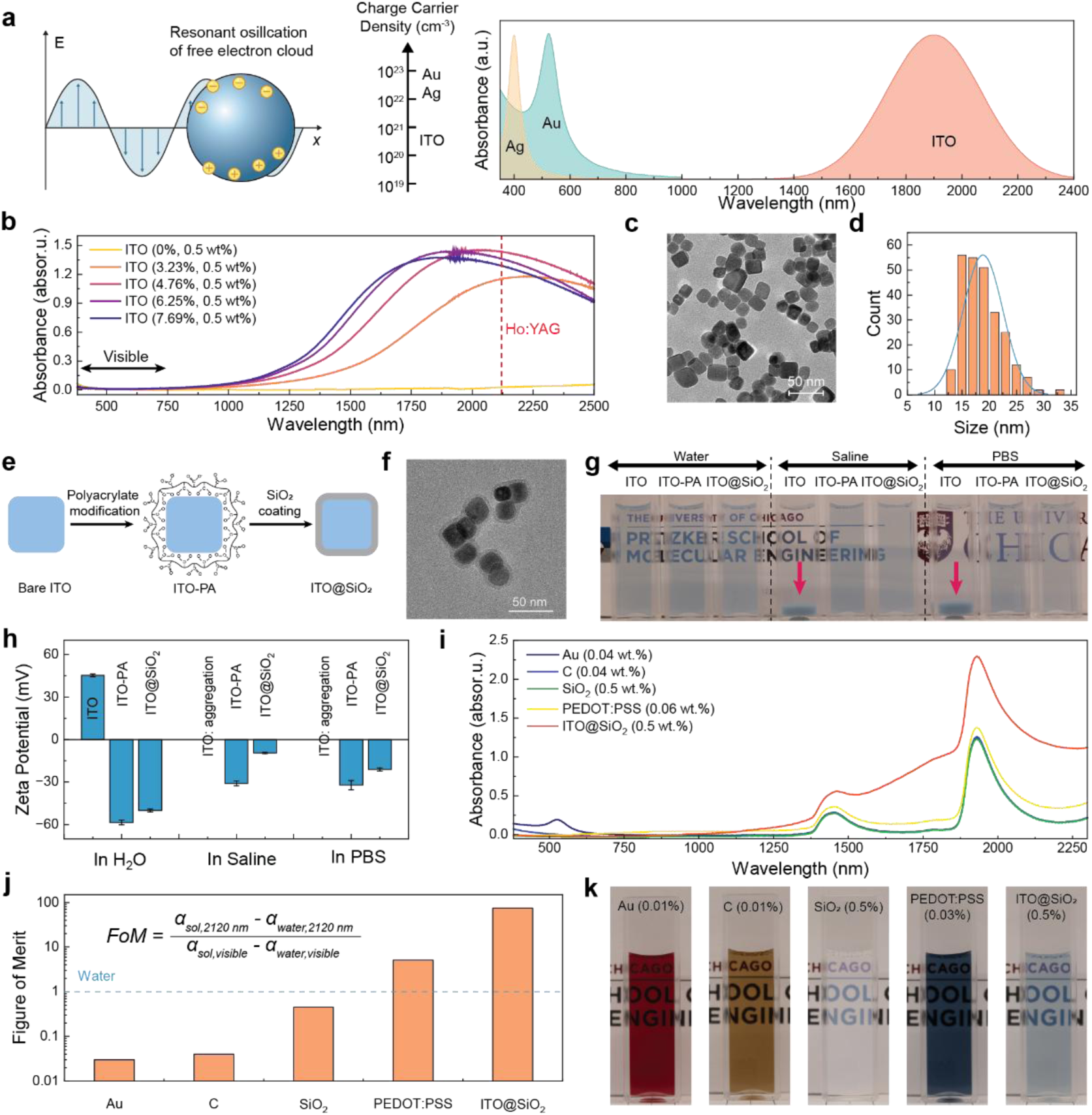
Optical and structural characterization of ITO and ITO@SiO_2_ nanoparticles. (a) Schematic illustration of localized surface plasmon resonance (LSPR) of noble metal and doped semiconductor nanoparticles. (b) Vis-NIR absorption spectra of ITO nanoparticles synthesized at various feeding ratios (water baseline). (c, d) Transmission electron microscopy (TEM) image (c) and corresponding size distribution (d) of the synthesized ITO nanoparticles (4.76% feeding ratio). (e) Schematic illustrating the surface modification of ITO nanoparticles. (f) TEM image of the resulting ITO@SiO_2_ core-shell nanoparticles. (g, h) Digital photographs demonstrating colloidal stability (g) and corresponding zeta potential measurements (h) of the nanoparticles before and after surface modification. (i) Vis-NIR absorption spectra of various comparative nanoparticle solutions (air baseline). (j) Figures of merit (FoMs) calculated from the absorption spectra of the respective solutions. (k) Macroscopic digital photographs of the nanoparticle solutions (path length: 1 cm).

Compared to alternative methods that utilize nonpolar solvents,^24,25^ synthesizing ITO nanoparticles in ethylene glycol allows them to be easily dispersed in water, yielding a highly transparent and homogeneous solution. Transmission electron microscopy (TEM) images reveal that ITO nanoparticles synthesized with a tin chloride feeding ratio of 4.76% possess a cuboidal shape with an average size of 18 nm (**Fig. 2c** and **2d**). While altering this feeding ratio during synthesis changes their absorption spectra, the nanoparticle morphology remains largely unchanged (**Supplementary Fig. 1**). However, because no surface ligands are introduced during synthesis, the bare ITO nanoparticles rely solely on electrostatic repulsion for stabilization in water. Consequently, their colloidal stability is highly sensitive to the ionic strength of the surrounding medium. As shown in **Fig. 2g**, when water is replaced with saline or phosphate-buffered saline (PBS), the bare ITO nanoparticles rapidly precipitate out of the solution, forming a bluish precipitate at the bottom of the cuvette (indicated by red arrows). To enhance their resistance to aggregation, we propose a two-step surface modification strategy (**Fig. 2e**). First, short polyacrylate (PA) ligands were grafted onto the ITO surface via electrostatic interactions between the negatively charged-COO⁻ groups and the positively charged ITO nanoparticles (**Fig. 2h**). A shift in zeta potential from positive to negative confirmed the successful grafting of these ligands (yielding ITO-PA). While these ITO-PA nanoparticles remained well-dispersed in both saline and PBS for several hours—demonstrating significantly improved stability over bare ITO—this stabilization proved insufficient under more extreme conditions. Specifically, when the ionic strength was further elevated by the addition of BegoStone, the ITO-PA solution gradually became turbid as the nanoparticles precipitated (**Supplementary Fig. 2**). The second step of our modification involves encapsulating the ITO-PA nanoparticles within a thin silica (SiO_2_) shell using a modified Stöber method. TEM confirmed the successful formation of an approximately 4 nm thick SiO_2_ shell (**Fig. 2f**). Protected by this robust inorganic layer, the nanofluid remained highly stable in both saline and PBS for several days, even in the presence of BegoStone. Such exceptional colloidal stability against salt-induced aggregation is a critical prerequisite for its practical application as a reliable irrigation fluid during LL.

To comprehensively evaluate the efficacy of ITO@SiO_2_ as a surrounding fluid in LL, we compared it against four alternative nanoparticles with distinct optical properties: gold (Au), carbon (C), silica (SiO_2_), and poly(3,4ethylenedioxythiophene) polystyrene sulfonate (PEDOT:PSS). We first confirmed their respective absorptive behaviors using a Vis-NIR spectrometer (**Fig. 2i**). Specifically, Au nanoparticles absorb strongly at ∼530 nm due to localized surface plasmon resonance; C nanoparticles exhibit broad-band absorption across the entire visible spectrum; SiO_2_ nanoparticles barely absorb light in either the visible or NIR regions; whereas PEDOT:PSS demonstrates moderate absorption spanning both ranges. To quantitatively assess each nanoparticle’s ability to absorb laser energy while maintaining visible clarity, we defined a figure of merit (FoM) as follows:

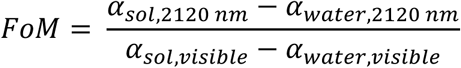

where α_*sol*,2120 *nm*_ and α_*water,*2120 *nm*_ represent the absorption coefficients of the solution and water at 2120 nm, respectively, while α_*sol,visible*_ and α_*water,visible*_ denote their average absorption coefficients in the visible range (380–750 nm). By definition, the FoM of pure water is set to 1. As shown in **Fig. 2j**, the FoMs of both Au and C fall far below 1 due to their strong absorption in the visible spectrum. The FoM of SiO_2_ is slightly below 1, primarily caused by visible transmittance loss from light scattering. In contrast, both PEDOT:PSS and ITO@SiO_2_ achieve an FoM greater than 1. Notably, the FoM of ITO@SiO_2_ is more than an order of magnitude higher than that of PEDOT:PSS, underscoring its exceptional near-infrared absorption coupled with negligible visible light attenuation.

The visual appearances of the nanoparticle solutions at specific concentrations, along with the BegoStones immersed in these corresponding fluids, are presented in **Fig. 2k** and **Supplementary Fig. 5, 6**. To objectively evaluate this visibility, we quantitatively calculated the relative luminance contrast between the BegoStone pixels and their surrounding background. Although the stone appears slightly blurred in the ITO@SiO_2_ solution, its boundaries remain clearly distinguishable, even at a relatively large camera-to-stone distance of 7 mm.

The efficacy of these various nanoparticle solutions as the surrounding fluid in LL was assessed using a spot treatment protocol (i.e., fixed-point laser irradiation) in a bench-top cuvette model (**Fig. 3a**). The laser parameters were set to a pulse energy (*E_p_*) of 0.4 J, a frequency (*f*) of 20 Hz under dusting mode, and a total pulse number (*PN*) of 60. As anticipated, the ablation efficiency (quantified by the ablated crater volume) was approximately proportional to the FoM of the nanomaterials (**Fig. 3b, c**). Across all tested standoff distances (SDs), the efficiency followed the order of ITO@SiO_2_ > PEDOT:PSS > water ≍ SiO_2_ > C ≍ Au. Remarkably, at an SD of 1 mm—a distance where LL in water produced minimal damage to the stone—LL in the ITO@SiO_2_ solution generated the most substantial damage, resulting in an extraordinary enhancement factor of 2660% (**Fig. 3d**). This finding clearly demonstrates that utilizing the ITO@SiO_2_ nanofluid can significantly expand the effective working distance of LL.

**Fig. 3.**
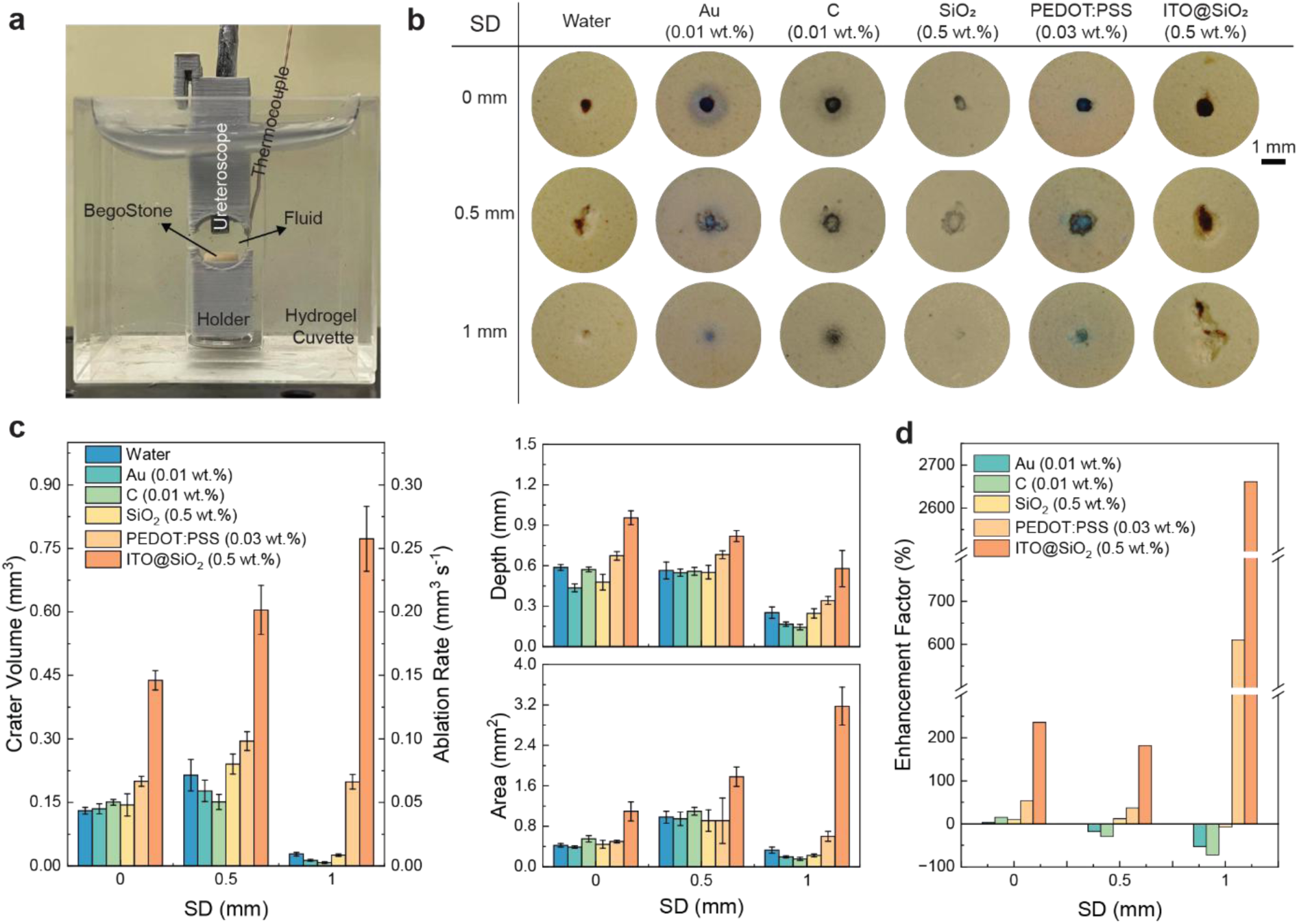
Ablation efficiency assessment in a bench-top cuvette model. (a) Digital photograph of the bench-top cuvette setup. (b) Digital photographs of the ablated craters produced in water and various nanoparticle solutions following spot treatment (*E_p_* = 0.4 J, *f* = 20 Hz, *PN* = 60). (c) Volume, depth, and profile area of the craters generated in the respective solutions. (d) Enhancement factors of the treatments in various solutions relative to water.

To elucidate the mechanisms driving this remarkable efficiency enhancement, we investigated bubble dynamics within the bulk fluids, as bubble expansion and collapse behaviors directly dictate the extent of stone damage. **Fig. 4a** illustrates the processes of bubble nucleation, expansion, and collapse in both water and the 0.5 wt.% ITO@SiO_2_ solution, while the corresponding quantitative bubble dimensions (length, width, and projected area) are detailed in **Fig. 4c**. Compared to water, bubbles generated in the ITO@SiO_2_ nanofluid exhibit three distinct differences: (1) vapor bubble nucleation occurs earlier following laser initiation; (2) the resulting bubble achieves significantly larger maximum dimensions in both length and width; and (3) bubble collapse occurs at ∼460 µs, which is extended by ∼110 µs compared to that in water (laser output power in **Fig. 4b**). These altered dynamics are a direct consequence of the ITO@SiO_2_ solution’s high absorption coefficient, which allows more laser energy to be absorbed and converted into heat for fluid vaporization. Consequently, the cavitation bubble in the nanofluid not only nucleates faster but also expands to a substantially larger maximum volume. Upon collapse, this larger bubble exerts a dramatically stronger mechanical impact on the stone surface—a conclusion directly supported by the two-fold increase in peak acoustic pressure measured via a hydrophone (**Fig. 4d**).

**Fig. 4.**
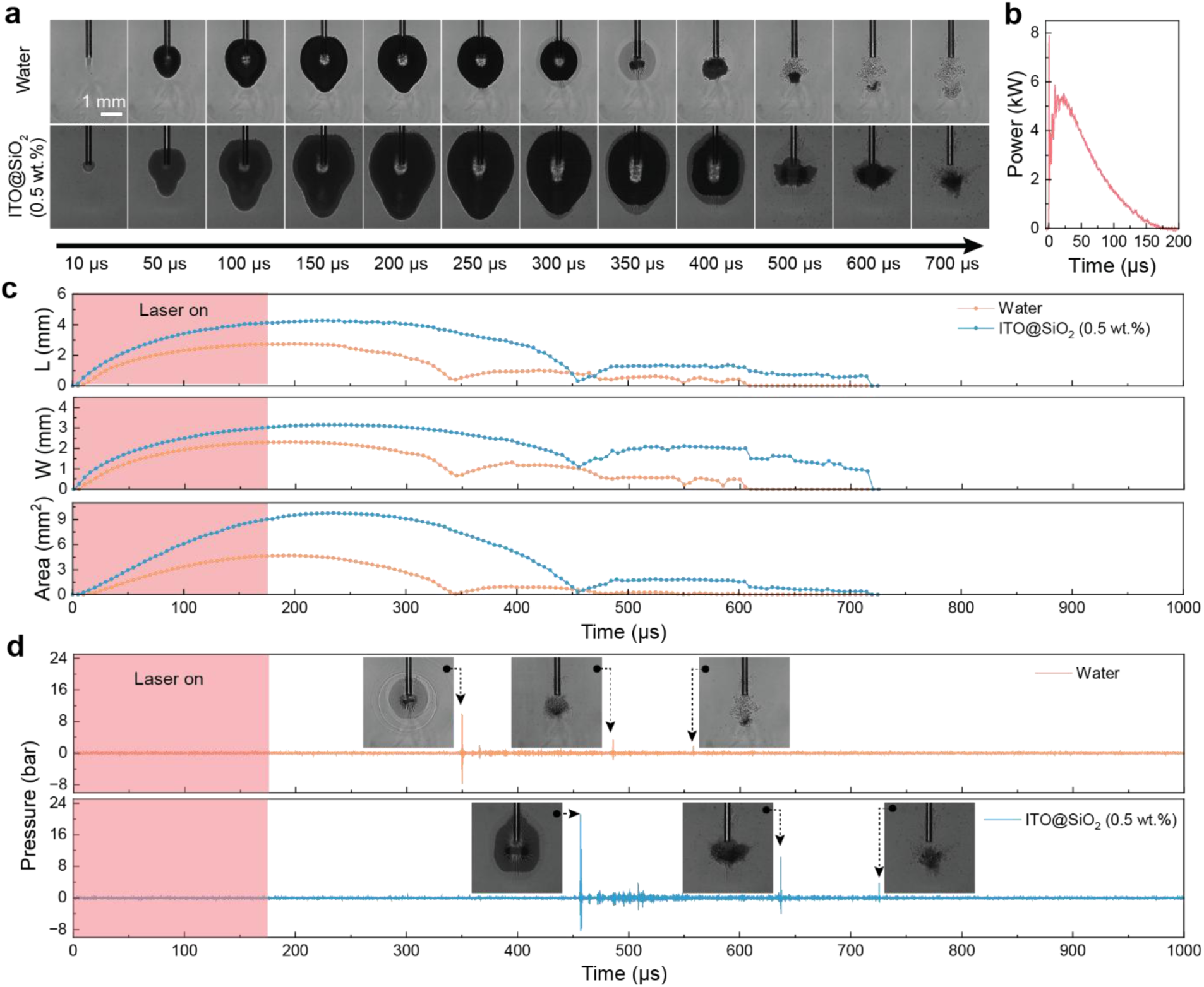
Bubble dynamics in bulk fluids. (a) High-speed imaging of bubble nucleation, expansion, and collapse triggered by a single laser pulse (*E_p_* = 0.4 J) in water (top) and 0.5 wt.% ITO@SiO_2_ solution (bottom). (b) Temporal profile of the laser output power (*E_p_* = 0.4 J). (c) Temporal evolution of the bubble length, width, and projected area corresponding to the events in (a). (d) Measured acoustic pressure changes during bubble expansion and collapse. Insets display the corresponding bubble morphologies at specific time points.

Beyond the enhanced cavitation damage, we hypothesize that improved photothermal ablation also contributes to the observed efficiency enhancement. This is primarily driven by the stone absorbing more energy as a result of faster bubble expansion. To mimic the influence of the stone’s presence on fluid dynamics, we deliberately controlled the SD between the fiber tip and a glass substrate. **Fig. 5a** illustrates bubble expansion in water and 0.5 wt.% ITO@SiO_2_ at an SD of 1 mm, alongside measurements of the fiber-to-apex distances (**Fig. 5b**, top). Notably, the bubble in the ITO@SiO_2_ solution reaches the substrate in ∼25 µs, which is 25 µs earlier than in water. This accelerated expansion allows a greater amount of laser energy to transmit through the vapor gap between the fiber tip and the stone, a phenomenon reflected by the enclosed area under the curves (**Fig. 5b**, bottom). The bubble behaviors at other SDs are summarized in **Supplementary Fig. 8**.

**Fig. 5.**
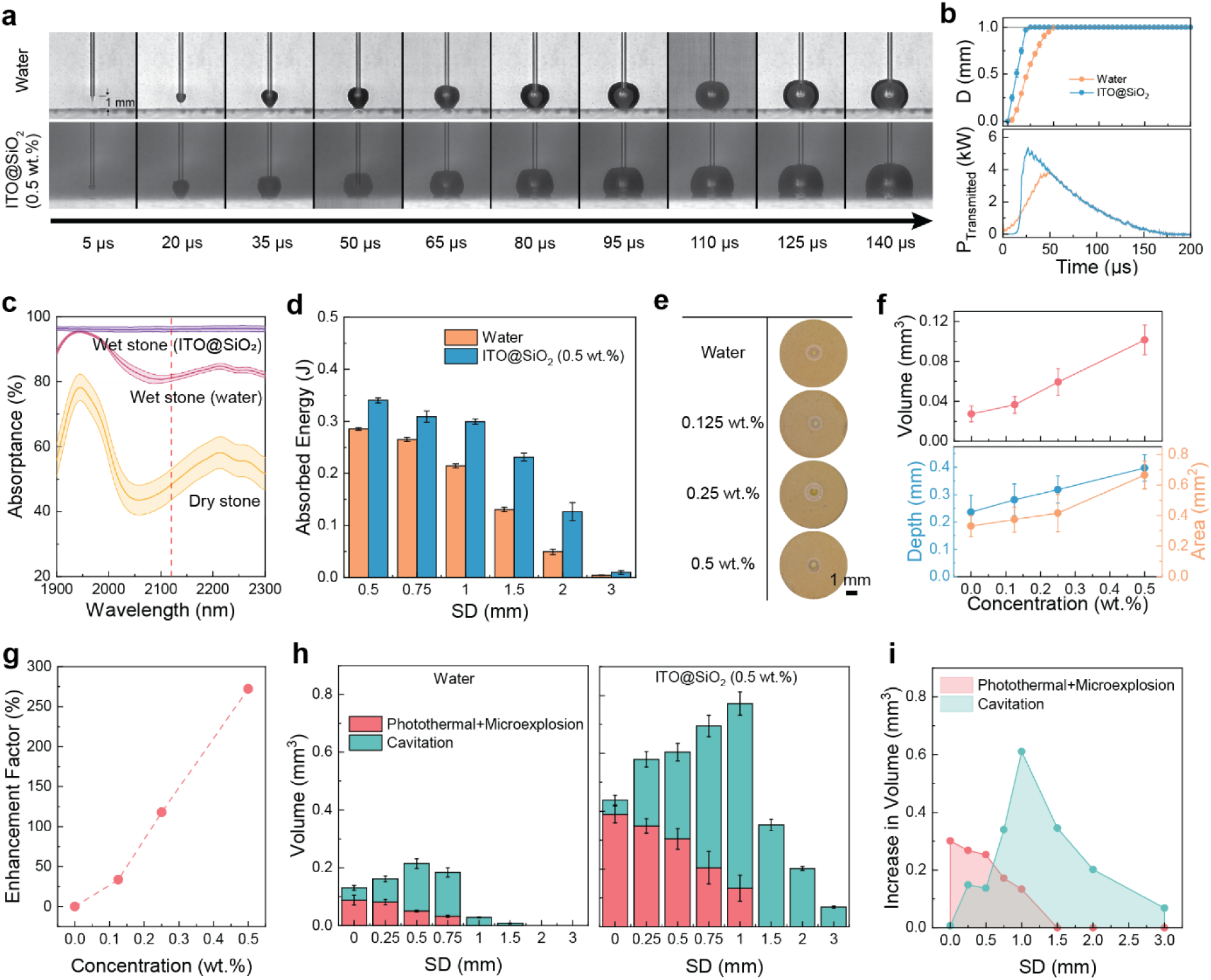
Decoupling the mechanisms of nanofluid-enhanced laser lithotripsy. (a) High-speed imaging of bubble expansion in front of a glass substrate at a standoff distance (SD) of 1 mm in water and the 0.5 wt.% ITO@SiO_2_ solution (*E_p_* = 0.4 J). (b) Measured apex-to-fiber distance (top) and calculated laser power transmitted through the glass substrate (bottom) at an SD of 1 mm. (c) Optical absorptance of dry, water-wetted, and nanofluid-wetted (0.5 wt.%) BegoStones. (d) Calculated theoretical energy absorbed by the BegoStones. (e) Digital photographs of ablated craters generated via LL on wetted BegoStones in air (SD = 0 mm). (f) Crater dimensions from panel (e), quantified via optical coherence tomography (OCT). (g) Ablation efficiency enhancement factors for LL performed on wetted BegoStones in air. (h) Decoupled quantitative contributions of photothermal/microexplosion effects versus cavitation damage to total stone ablation across various SDs. (i) Additional ablation damage achieved in the 0.5 wt.% ITO@SiO_2_ solution relative to water.

Alongside the transmitted energy, the absorptance of the stone must also be considered, as it determines how efficiently the laser energy is utilized. It is well established that BegoStone contains numerous micropores into which fluid can diffuse, thereby increasing its absorption of laser energy. Specifically, a dry stone absorbs only 48% of the incident laser energy, whereas a stone wetted with water absorbs 81%. Remarkably, when the BegoStone is wetted with the 0.5 wt.% ITO@SiO_2_ nanofluid, its laser energy absorptance peaks at 96% (**Fig. 5c**). Such heightened absorptance facilitates superior light-to-heat conversion, driving both photothermal ablation and internal microexplosions.

By taking both the transmitted energy and the stone’s absorptance into account, the theoretical energy absorbed by the stone can be calculated (**Fig. 5d**). The results clearly demonstrate that at every tested SD, BegoStone immersed in ITO@SiO_2_ absorbs a significantly greater amount of laser energy. This surplus energy is converted into heat, which melts the stone (photothermal ablation) and induces internal pressure gradients (microexplosions). To validate our hypothesis and explicitly isolate this effect from cavitation damage, we performed LL on wetted stones in air (**Fig. 5e**-**g**). The resulting ablated crater volume is roughly proportional to the ITO@SiO_2_ concentration and, consequently, the fluid’s absorption coefficient. The most extensive damage was achieved using 0.5 wt.% ITO@SiO_2_—the highest concentration investigated—yielding a remarkable enhancement factor of 272%.

The preceding experiments illustrate that photothermal ablation, cavitation damage, and potentially internal microexplosions all contribute to the remarkable enhancement in the ablation efficiency of ITO@SiO_2_. However, the quantitative contribution of each mechanism remains unclear. To address this, we conducted an additional spot treatment experiment in the bench-top cuvette model (*E_p_* = 0.4 J, *f* = 20 Hz, *PN* = 60) by strategically reducing the distance between the ureteroscope and the fiber tip. As previously established, this configuration causes the cavitation bubble to collapse against the ureteroscope rather than the stone substrate, thereby effectively eliminating the cavitation-induced damage (**Extended Data Fig. 1a-b**). Consequently, the crater volumes measured in this specific setup represent the combined effects of solely photothermal ablation and microexplosions. By subtracting these volumes from those obtained under standard LL settings, we can precisely isolate and quantify the specific contribution of the cavitation-induced damage (**Fig. 5h**).

The results indicate that the efficiency of photothermal ablation and microexplosions gradually decreases as the SD increases in both water and ITO@SiO_2_. This observation is consistent with our calculations showing reduced absorbed energy at larger SDs (**Fig. 5d**). Consequently, to maximize photothermal- and microexplosion-induced damage, the fiber tip must be positioned as close to the stone surface as possible. Conversely, cavitation-induced damage does not vary monotonically with SD. Instead, there exists an optimal SD that maximizes the cavitation effect: 0.5 mm in water and 1 mm in the ITO@SiO_2_ solution. Notably, across all three mechanisms—photothermal ablation, microexplosions, and cavitation—damage was significantly enhanced when the irrigation fluid was switched from water to ITO@SiO_2_ (**Fig. 5i**). This suggests that to fully exploit the benefits of the ITO@SiO_2_ nanofluid, LL should be performed at an SD where all ablation mechanisms are synergistically optimized (i.e., 1 mm in this specific spot treatment setup). Furthermore, we observed that this optimal SD is highly dependent on the laser treatment parameters (**Extended Data Fig. 2a, b**). For instance, when the *PN* was reduced to 10 while keeping *E_p_* and *f* constant, the optimal SD in ITO@SiO_2_ decreased to 0.5 mm. This shift occurs likely because cavitation damage relies on a threshold-dependent cumulative effect, requiring a critical number of pulses to inflict maximum structural damage.

The remarkable improvement in ablation efficiency achieved with the ITO@SiO_2_ nanofluid is not limited to a single power setting. Experimental results demonstrate that across all investigated power settings (from 2 W to 8 W), treatments in the ITO@SiO_2_ nanofluid exhibited a >200% enhancement compared to those in water (*E_p_* = 0.4 J, *f* = 20 Hz, *PN* = 60, SD = 0 mm; **Extended Data Fig. 2c**). Furthermore, while spot treatments are useful for mechanistic studies, scanning treatments—where the laser fiber is continuously moved across the stone’s surface—more accurately reflect routine clinical practice. To evaluate the nanofluid’s performance under these clinically relevant conditions, we utilized a motorized stage to control the fiber’s moving speed over a BegoStone slab. We investigated two scan speeds and three SDs with a 2-min lasing duration (**Extended Data Fig. 2d, e**). Factoring in the fiber diameter, a scan speed of 0.2 mm s^-1^ roughly equates to 50 pulses per fixed spot, whereas 1 mm s^-1^ equates to 10 pulses. Consistent with our spot treatment findings, the slower scan speed yielded a more pronounced efficiency enhancement (>100% across all SDs), presumably due to the cumulative nature of cavitation damage. Interestingly, although the faster scan speed (1 mm s^-1^) resulted in a relatively smaller enhancement factor (∼50%), it achieved a higher overall ablation rate than the slow scan speed. This occurs because a faster scan continuously exposes fresh stone surface to the laser, ensuring that energy is efficiently utilized to ablate the superficial material at the designated SD, rather than being attenuated at the bottom of a deepening crater.

To evaluate whether ITO@SiO_2_ nanoparticle-assisted irrigation alters thermal burden during LL, we utilized an anatomically accurate kidney phantom model using ballistic hydrogel. During the treatment, seven thermocouples were positioned throughout the upper calyx containing the BegoStone to monitor the temperature at the fluid-gel interface, thereby mimicking the tissue boundary temperature (**Fig. 6a**). LL was performed at 0.4 J and 20 Hz using a 50% duty cycle (i.e., 30 s on/30 s off), corresponding to a total active lasing duration of 3 min. Continuous room-temperature irrigation consisting of either saline or 0.5 wt.% ITO@SiO_2_ dispersed in saline was maintained at a flow rate of 20 mL min^-1^ throughout the procedure (**Extended Data Fig. 3a**).

**Fig. 6.**
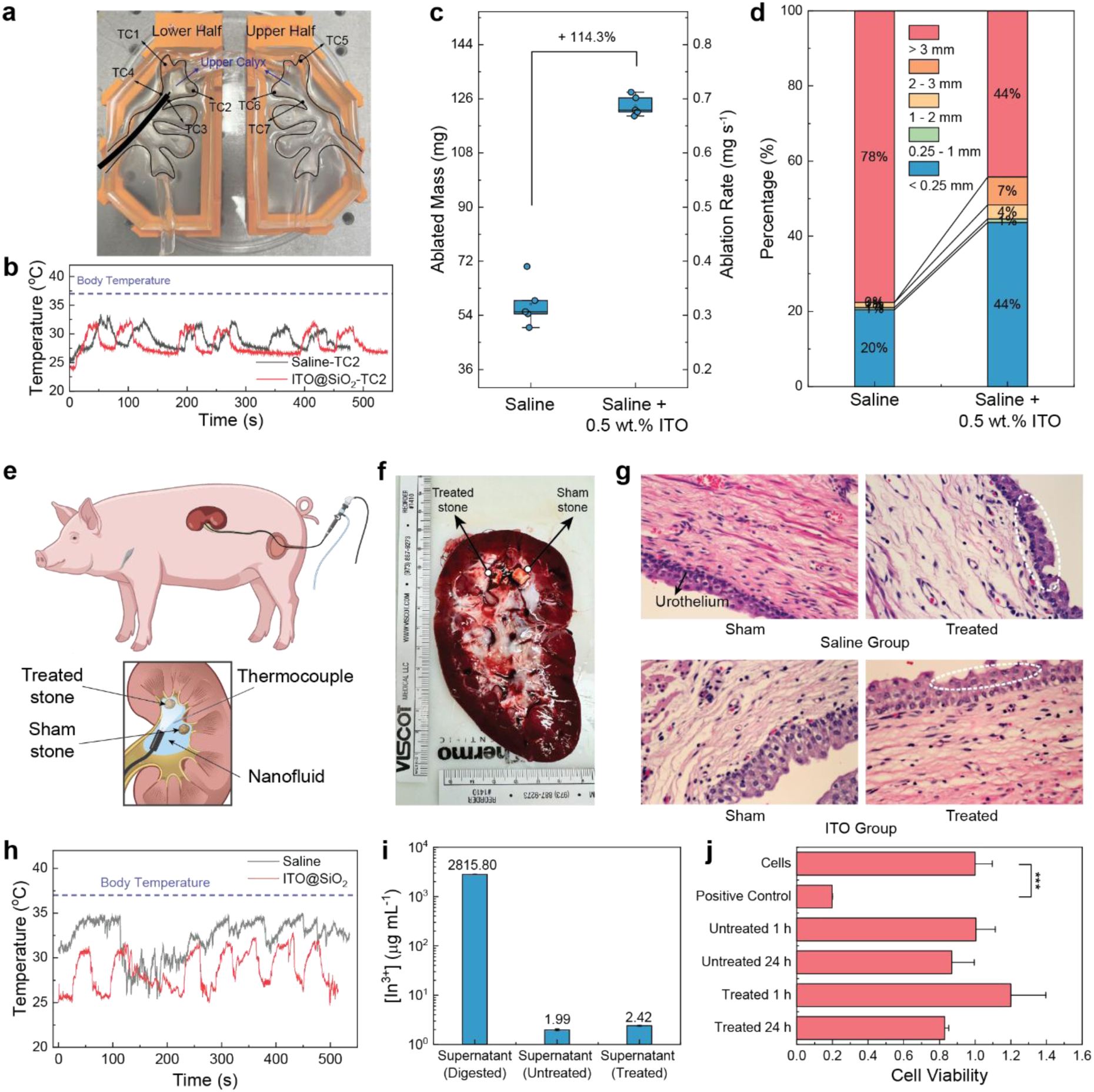
Safety and efficacy assessments of nanofluid-assisted laser lithotripsy. (a) Digital photograph of the ballistic hydrogel kidney phantom model, used for thermal mapping during laser lithotripsy (LL), showing the locations of the seven thermocouples positioned around the upper calyx. (b) Representative temperature profiles recorded in the upper calyx during LL performed using saline or 0.5 wt.% ITO@SiO_2_ irrigation. The dashed line indicates normal body temperature. (c) Total ablated stone mass and corresponding ablation rates (calculated based on active lasing time) for BegoStones treated in the hydrogel kidney model. (d) Size distribution of the collected stone fragments post-treatment. (e) Schematic illustration of the in vivo LL in pig. (f) Digital photograph of the explanted porcine kidney following the in vivo experiments. (g) Representative hematoxylin and eosin (H&E)-stained histological sections of the renal urothelium following sham or laser treatment using saline or ITO@SiO_2_-assisted irrigation. Dashed circles indicate mild superficial urothelial alterations. (h) In vivo intrarenal temperature profiles recorded during LL in the porcine model. The dashed line indicates normal body temperature. (i) Concentrations of In^3+^ in the supernatants of the untreated and laser treated ITO@SiO_2_ (0.5 wt.%). (j) Viability of mIMCD-3 cells, measured with a MTT assay, after incubation with untreated and laser treated ITO@SiO_2_ (0.5 wt.%) for 1 and 24 h. Exposure to triton X-100 (10% in PBS, 30 s) was used as a positive control to decrease cell viability. Significance was measured using 1-way ANOVA with Dunnett’s multiple comparisons post hoc. ***p <0.001, not significant comparisons are not presented.

Under both irrigation conditions, fluid temperature transiently increased during laser activation before reaching a steady-state plateau. Importantly, the maximum sustained temperature recorded across all seven thermocouples remained below 34 °C (**Fig. 6b** and **Extended Data Fig. 3b**), substantially lower than temperatures associated with thermally induced renal injury (> 43 °C). Notably, the temperature evolution observed in the ITO@SiO_2_ nanofluid closely mirrors with that of saline alone, suggesting that enhanced optical absorption by the nanofluid does not translate into measurable thermal accumulation under clinically relevant irrigation conditions.

Following treatment, the stone fragments were collected, dried, and size-fractionated using meshed sieves. Relative to saline only irrigation, treatment using the ITO@SiO_2_ nanofluid yielded a 114% improvement in ablation efficiency. Moreover, fragment analysis revealed a marked increase in the proportion of dust-sized particles (≤0.25 mm), indicating more efficient stone dusting in the nanofluid-assisted group (Note: Fragments larger than 3 mm predominantly corresponded to intentionally untreated residual stone because the procedures were terminated prior to complete stone clearance; **Fig. 6c** and **6d**).

To investigate the translational applicability and renal tissue safety of ITO@SiO_2_ nanofluid irrigation, in-vivo LL experiments were performed in anesthetized porcine kidneys (**Fig. 6e**). Two BegoStones were retrogradely implanted into the upper calyx of each kidney.^26^ One stone was treated with LL while the second served as a sham control exposed only to irrigation to account for mechanical injury (**Fig. 6f**). A single thermocouple was placed at the urothelial junction within the calyx of interest containing the stone. A total of two pigs (four kidneys) were included in this vivo study. Identical laser settings and duty cycles utilized in the hydrogel studies were performed in each renal unit, using either saline (N=2) or ITO@SiO_2_/saline irrigation (N=2). Following the procedure, bilateral kidneys were explanted for histopathological examination, and large stone fragments were extracted to calculate ablation efficiency.

Histological examination of hematoxylin and eosin (H&E)-stained tissue sections was performed. The sham kidneys revealed intact urothelial morphology with cohesive cellular arrangements following exposure to either saline or ITO@SiO_2_/saline irrigation alone (**Fig. 6g**). In the laser-treated kidneys, mild superficial urothelial alterations including subepithelial clefting in the saline-treated group and focal apical surface disruption in the ITO-treated group (highlighted by dashed circles), were observed. These limited superficial changes are likely attributable to mechanical injury from ureteroscopic stone manipulation. Additionally, focal hemorrhage was observed in the second pig with localized disruption of superficial microvasculature (**Supplementary Fig. 10**). Importantly, no evidence of deep thermal coagulation, tubular necrosis, or structural injury to the renal parenchyma was identified.

Glomeruli and renal tubules remained histologically intact in both treatment groups. Consistent with observations from the hydrogel model, intrarenal fluid temperatures remained below 35 °C throughout the procedure without sustained hyperthermia, thereby confirming the thermal safety of nanoparticle-assisted irrigation (**Fig. 6h**). In terms of in vivo efficiency, the ITO@SiO_2_/saline group achieved an average ablated mass of 112 mg and an ablation rate of 0.62 mg s^-1^, representing a 28% enhancement over standard saline irrigation (87.8 mg and 0.488 mg s^-1^, respectively).

The localized high temperatures generated during LL could potentially induce the dissolution of ITO, thereby elevating the concentration of toxic In^3+^ in the surrounding fluid. To evaluate this risk, we quantified the In^3+^ concentration in the supernatant using inductively coupled plasma mass spectrometry (ICP-MS, **Fig. 6i**). Following a 1-min laser treatment, the In^3+^ concentration increased only marginally from 1.99 µg mL^-1^ to 2.42 µg mL^-1^. This minimal change indicates that the localized heating during LL causes negligible dissolution of the ITO nanoparticles. For comparison, the total In^3+^ concentration in the supernatant reached ∼2816 µg/mL when the ITO particles were completely digested in HNO_3_. Furthermore, the in vitro cytotoxicity of the nanofluid was evaluated using murine inner medullary collecting duct (mIMCD-3) epithelial cells (**Fig. 6j**). Cell viability remained uncompromised even following a 24-h incubation with either the untreated or laser-treated ITO@SiO_2_ nanofluids. This excellent short-term biocompatibility, primarily attributed to the protective SiO_2_ shell, further validates the safety and translational potential of the nanofluid in LL.

Because ITO exhibits a broad NIR absorption band, this nanofluid strategy should theoretically benefit other emerging laser technologies, including the TFL (1940 nm) and Tm:YAG laser (2010 nm). Validating this, spot treatments using 0.5 wt.% ITO@SiO_2_ yielded efficiency enhancements exceeding 130% for the TFL (*E_p_* = 0.2 J, *f* = 20 Hz, *PN* = 60) and 240% for the Tm:YAG laser (*E_p_* = 0.2 J, *f* = 25 Hz, *PN* = 60) (**Extended Data Fig. 4**). Ultimately, this demonstrates the versatile potential of engineered nanofluids to augment ablation efficiency across the entire spectrum of NIR lithotripsy lasers. However, optimizing this nanofluid specifically for other laser systems remains an important avenue for future research.

## Discussion

In this study, we introduce a novel nanofluid irrigation strategy designed to enhance the treatment efficiency of LL using the widely adopted clinical gold-standard: Ho:YAG laser. Compared to previous nanoparticle-assisted LL strategies, our approach offers two distinct translational advantages. First, the NIR absorption peak of the ITO@SiO_2_ nanoparticles precisely matches the wavelength of the Ho:YAG laser, avoiding the reliance on experimental or less common lasers operating in the 500–1500 nm range. Second, our method bypasses the need for complex stone pretreatment. By simply replacing standard saline irrigation with the engineered nanofluid, this approach integrates seamlessly into established clinical workflows without imposing any additional learning curve on urologists.

We previously demonstrated the feasibility of using a PEDOT:PSS nanofluid to improve Ho:YAG LL efficiency. However, while PEDOT:PSS exhibits NIR absorption, its moderate absorption in the visible spectrum restricts the maximum usable concentration, ultimately capping the fluid’s absorption coefficient increase at only 25%. In contrast, the negligible visible-light absorption of the ITO@SiO_2_ nanoparticles developed in this work permits the use of a substantially higher concentration (up to 0.5 wt.%) without compromising ureteroscopic visibility. This breakthrough leads to a remarkable 395% increase in the absorption coefficient at 2120 nm, which fundamentally drives the superior ablation outcomes observed herein.

Benefiting from this remarkably high absorption coefficient, LL in the ITO@SiO_2_ nanofluid achieves an ablation rate of 0.25 mm^3^ s^-1^ at the optimal SD during spot treatments. This significantly surpasses the maximum ablation rate of 0.075 mm^3^ s^-1^ observed in water under the same experimental setup. Analysis of bubble dynamics reveals that this increased absorption coefficient drives both faster bubble expansion and a larger maximum bubble size. Both characteristics critically contribute to the improved ablation efficiency: (1) The accelerated bubble expansion allows additional laser energy to transmit through the vapor gap between the fiber tip and the stone. Concurrently, the infiltration of ITO@SiO_2_ nanoparticles into the porous stone matrix increases the stone’s direct absorption of this laser energy, ensuring highly efficient light-to-heat conversion. This compounding effect drastically improves photothermal ablation and internal microexplosions. (2) The collapse of the resulting larger bubble generates a two-fold increase in acoustic pressure, inflicting significantly stronger cavitation damage on the stone. Furthermore, our quantitative decoupling analysis indicates that enhanced photothermal ablation and microexplosions dominate at shorter SDs (0–0.5 mm), whereas cavitation damage becomes the primary contributor at larger SDs (0.5–3.0 mm).

The thermal and biological safety profile of this strategy was systematically evaluated using both hydrogel kidney phantoms and in vivo porcine models under clinically relevant irrigation conditions. To the best of our knowledge, this study represents the first in vivo demonstration of nanoparticle-enhanced LL in a large-animal model. Despite the substantially enhanced optical absorption of the ITO@SiO_2_ nanofluid, intrarenal temperatures remained below 35 °C and closely matched those observed with saline irrigation alone. This suggests that improved laser–fluid coupling does not result in measurable thermal accumulation. This finding is particularly important because thermal injury remains a major concern during high-power endoscopic lithotripsy, especially as higher laser energies and pulse frequencies are adopted to enhance stone fragmentation efficiency. Our findings suggest that engineering the optical properties of the irrigation medium provides an alternative route to improving lithotripsy efficiency without escalating laser power output. Rather than serving solely as a passive cooling medium, the irrigation fluid can instead be engineered as an active optical component of the lithotripsy system, enhancing local laser energy utilization while continuous irrigation dissipates excess heat from the collecting system, thereby decoupling improved ablation efficiency from thermal accumulation. Importantly, no evidence of deep thermal injury or renal parenchymal damage was observed following nanoparticle-assisted LL, suggesting that enhancement of laser-induced cavitation and photothermal effects can remain spatially confined to the immediate stone environment under clinically relevant irrigation conditions.

Additionally, in vitro cytotoxicity assays—conducted by incubating mIMCD-3 cells with the 0.5 wt.% ITO@SiO_2_ nanofluid—showed no reduction in cell viability even after 24 hours of exposure. This strongly underscores the excellent acute biocompatibility of the engineered ITO@SiO_2_ nanoparticles.

Although our surgical protocol involves flushing the kidney with normal saline at the conclusion of the procedure to remove the ITO@SiO_2_ nanofluid, guaranteeing the complete clearance of all nanoparticles remains clinically challenging. Consequently, the potential accumulation, systemic biodistribution, and long-term in vivo toxicity of ITO@SiO_2_ nanoparticles within the urinary tract and systemically require comprehensive future investigation. Furthermore, the clinical efficacy of this approach must be validated using actual human urinary calculi. Given their high compositional heterogeneity, clinical stone samples may behave differently than standardized BegoStone phantoms—particularly regarding their inherent porosity and optical properties—which could ultimately influence the coupled ablation mechanisms.

### Experimental Section

#### Materials

The following chemicals were used as received without further purification. Indium chloride tetrahydrate (InCl_3_•4H_2_O, 97%), tin chloride pentahydrate (SnCl_4_•5H_2_O, 98%), sodium hydroxide (NaOH, 97+%) polyacrylic sodium salt (average Mw 2.1k), anhydrous ethanol (99.5+%, 200 proof), ammonium hydroxide solution (28-30% NH_3_ basis), tetraethyl orthosilicate (TEOS, 98%), l-ascorbic acid (99%), polyvinylpyrrolidone (PVP, average Mw 40k), Gold(III) chloride trihydrate (HAuCl_4_, 99.9+%), sodium borohydride (NaBH_4_, 98+%), sodium citrate tribasic dihydrate (99+%), potassium iodide (99+%), cellulose membrane (molecular weight cut-off: 14k), phosphate buffered saline (PBS, pH 7.2) were purchased from Sigma-Aldrich. Aqueous dispersion of PEDOT:PSS conductive polymer (Clevios PH 1000) was purchased from MSE Supplies. Ethylene glycol (99+%) was purchased from Thermo Fisher Scientific.

#### Synthesis of ITO nanoparticles

The solvothermal reaction for ITO nanoparticles synthesis was adopted from the literature with slight modification.^27^ Typically, to synthesize ITO nanoparticles with a feeding ratio of 4.76%, 2.932 g InCl_3_•4H_2_O and 0.175 g SnCl_4_•5H_2_O were dissolved in 20 mL ethylene glycol to form solution A. To obtain solution B, 1.2 g NaOH was dissolved in 20 mL ethylene glycol with the help of sonication. Solution B was added to solution A dropwise under stirring and after stirring the mixture at room temperature for 1 h, the mixture was transferred to a 100 mL PPL lined hydrothermal reactor. The reaction was heated to 250 °C and allowed to react at this temperature for 24 h. After the reaction naturally cooled down to RT, the supernatant was decanted, and the precipitates were purified with EG twice and water twice to remove the unreacted residues. Centrifugation and sonication were used in between the purification cycles. The final product was dispersed in water, and its concentration was determined using the oven-drying method.

#### Synthesis of ITO@SiO_2_ nanoparticles

ITO nanoparticles were first grafted with polyacrylate to improve their colloidal stability prior to the silica coating. 0.54 g polyacrylic sodium salt was dissolved in 96 mL water and 12 mL 3 wt.% ITO solution was added. The mixture was allowed to react in the sonication bath for 1 h. The extra ligands were separated from the ITO nanoparticles by centrifugation, and the ITO nanoparticles were dispersed in ∼10 mL water. For the silica coating, 72 mL EtOH, 1 mL NH_4_OH and 0.5 mL TEOS were added to the polyacrylate modified ITO solution. The mixture was stirred at RT for 2 h before it was transferred to a dialysis tubing cellulose membrane (molecular weight cut-off:14000) for dialysis in EtOH. After dialyzing in 700 mL EtOH for over 24 h, the ITO@SiO_2_ nanoparticles were separated from EtOH using centrifugation and finally dispersed in water.

#### Transmission electron microscopy (TEM)

The NP solution was dropped onto a copper grid and removed the water after 30 min. TEM images were taken on the FEI Tecnai G2 F30 300kV Super Twin Electron Microscope equipped with a CCD camera.

#### Dynamic light scattering and zeta potential

The hydrodynamic size and zeta potential of the dilute solutions of ITO, ITO-PA and ITO@SiO_2_ were measured on Malvern Zetasizer Nano ZS.

#### UV-VIS-NIR absorption spectrum measurement

The UV-VIS-NIR absorbance spectra of solutions were measured using a UV–vis–NIR spectrophotometer (Cary 5000, Agilent, Santa Clara, CA, USA). During the measurement, the solution was filled in a cuvette with a path length of 0.2 mm (IR quartz, FireflySci, Inc. Northport, NY, USA), and the scan range was set to be 380–2500 nm (resolution: 1 nm and scan rate: 600 nm min^−1^). The slit width was 1 nm. An empty cuvette was used as the baseline when collecting data.

The visible transmission spectra were measured on the same spectrophotometer with the scan range of 380–750 nm. A glass cuvette with a path length of 10 mm filled with water was used as the baseline when collecting data. The average transmittance of a solution in visible is calculated by taking the average value.

#### NIR diffusive reflectance spectrum measurement

The NIR diffusive reflectance of the BegoStone samples (25 mm in diameter and 7 mm in thickness) was measured using a UV–vis–NIR spectrometer (Shimadzu UV3600 Plus) equipped with an integrating sphere. The scan range was set to be 1950–2250 nm (resolution: 1 nm and scan rate: medium). The slit width was 1 nm. A standard white plate coated with BaSO_4_ was used as the reference. Stones of this size were chosen to ensure they cover the whole laser beam size (15 mm × 6 mm). Stones were dried at 80 °C overnight to remove the moisture, and they were immersed in the corresponding fluids for 5 min prior to the measurement. After being taken out of the fluids, stones were wiped with papers tissues to remove any liquid on their surfaces. For each fluid, the same measurement was repeated on five different stones with both sides to account for the variation in stones. Since BegoStone was very thick (7 mm), we assume no light could transmit through, so stone absorption = 1-reflectance.

#### In vitro stone phantom experiments

BegoStone phantoms (BEGO USA) were prepared at a 5:2 powder-to-water ratio following previously reported protocols and treated using a clinical Ho:YAG laser system (H Solvo 35 W, Dornier MedTech) with a 270 µm fiber (Dornier SingleFlex 270, NA = 0.26) operated in dusting mode. Two experimental setups were used. For spot treatments, 6 × 6 mm cylindrical stones were positioned in a quartz cuvette with hydrogel-lined walls. A 3D-printed insert containing a spherical chamber (radius = 5 mm) was filled with 2 mL of test fluid to mimic a single calyx environment. The ureteroscope, with a fiber tip advanced 5 mm beyond the scope end, was fixed on a 3D translational stage to precisely control the standoff distance (SD) between the fiber tip and the stone surface. For scanning treatments, larger BegoStone slabs (23 × 23 × 10 mm^3^) were immersed in ∼20 mL of fluid, and the fiber-integrated ureteroscope was mounted on a motorized stage to enable controlled translation across the stone surface at defined speeds and SDs. Laser irradiation was applied in pulse trains, typically 10 or 60 pulses for spot treatments and 2400 pulses for scanning treatment.

Different experimental configurations were implemented to evaluate ablation efficiency and isolate underlying mechanisms. First, to compare dusting efficiency across different nanofluids, spot treatments were performed in the cuvette setup filled with water, 0.01 wt.% Au, 0.01 wt.% C, 0.5 wt.% SiO_2_, 0.03 wt.% PEDOT:PSS, or 0.5 wt.% ITO@SiO_2_ at SDs of 0, 0.5, and 1.0 mm under the Ho:YAG laser settings of 0.4 J and 20 Hz (i.e., a laser power of 8 W). For the 0.5 wt.% ITO@SiO_2_ nanofluid, laser power was further varied from 2 W to 8 W by adjusting pulse energy and frequency in dusting mode (0.2 J/10 Hz, 0.2 J/20 Hz, 0.4 J/15 Hz, and 0.4 J/20 Hz). Crater damage was quantified using optical coherence tomography (OCT), from which crater volume, maximum depth and profile area were extracted.

Second, to illustrate the enhanced photothermal ablation and microexplosion effects by the ITO@SiO_2_ nanofluids, wetted-stone treatments were performed in air by applying 50 µL of ITO@SiO_2_ at concentrations of 0.125, 0.25, and 0.5 wt.%. to the stone surface and allowing it to fully absorb prior to laser irradiation. The fiber tip was placed in direct contact with the stone surface (i.e., SD = 0 mm) to maximize laser energy delivery. The resultant crater damage was then compared with that produced with pure water under the same conditions.

Third, to isolate cavitation-induced damage, the offset distance between the fiber tip and the scope end was shortened from 5 mm to 0.25 mm so that bubble collapse was distracted away from the stone surface. By comparing stone damage produced under the standard and shortened offset configurations, the contribution of cavitation to stone damage was quantified over a wider SD range of 0–3 mm in both water and 0.5 wt.% ITO@SiO_2_ nanofluids under 0.4 J and 20 Hz settings.

Furthermore, to assess clinical relevance, scanning treatments were performed at two representative scan speeds of 0.2 mm/s and 1 mm/s and three SDs of 0.25 mm, 0.5 mm and 1 mm. Ablation efficiency produced in both water and 0.5 wt.% ITO@SiO_2_ was quantified by mass loss after drying stones.

#### Bubble Dynamics

Bubble dynamics were visualized in water and 0.5 wt.% ITO@SiO_2_ nanofluid under Ho:YAG laser irradiation (0.4 J, 20 Hz) using an ultra-high-speed camera (Kirana-M5, Specialised Imaging) operated at 0.2 million frames per second. Backlighting was provided by a 10-ns pulsed laser illumination system (SI-LUX-640, Specialised Imaging). Simultaneously, acoustic emissions generated during bubble collapse were measured using a needle hydrophone (HNC-1000, Onda) connected to a digital oscilloscope (HDO-6104, Teledyne LeCroy). The hydrophone was positioned 10 mm from the fiber tip, and the measured pressure was normalized to a distance of 1 mm. To further quantify laser energy transmission during bubble expansion at different SDs, bubbles were generated in front of a 1-mm-thick glass slide and imaged under the same high-speed conditions.

#### In vitro stone phantom experiments

The hydrogel kidney model was prepared as previously described,^28^ with BegoStone phantoms placed in the upper calyx and treated by the Ho:YAG laser at 0.4 J and 20 Hz settings under a room-temperature irrigation of 20 mL min⁻¹, while fluid temperature in the calyx was monitored by seven K-type thermocouples (5SRTC-TT-K-36-36, Omega) at 0.2-s intervals. After treatment, the remaining stone fragments were collected, dried, and sieved to quantify treatment efficiency and fragment-size distribution produced in both water and 0.5 wt.% ITO. Fragments were grouped by size: <0.25 mm, 0.25–1 mm, 1–2 mm, 2–3 mm, and >3 mm.

#### LL in porcine kidney

Institutional Animal Care and Use Committee approval was obtained for this study. We performed stone implantation followed by ureteroscopy and LL on two live anesthetized female pigs (75–100 lbs). Each pig underwent sequential implantation and treatment of 4–6 × 10 mm pre-weighed cylindrical soft BegoStone phantoms(BEGO™, Lincoln, RI) in both kidneys. Stone phantoms were prepared with a 5:2 powder-to-water ratio and soaked in water for 24 h before implantation to achieve mechanical properties comparable to human kidney stones. Stone implantation was performed by urology residents, while ureteroscopy with LL was performed by an endourology fellow.

General anesthesia was administered. Cystoscopy and bilateral ureteral catheterization were performed followed by a laparotomy and proximal ureterotomy for stone implantation. After placing two stones in the kidney, the ureterotomy was closed using 4−0 absorbable suture, and watertight closure was confirmed. An 11/13 Fr ureteral access sheath was placed over a wire into the proximal ureter, and a single-use flexible ureteroscope (AXIS II, Dornier MedTech, Germany; outer diameter 8.7 Fr) was advanced through the sheath. This resulted in a sheath-to-scope ratio of 1.26. The two stones were manipulated into separate nonadjacent calyces. Using endoscopic guidance, a K-type thermocouple (5SRTC-TT-K-36-36, OMEGA, Norwalk, CT, USA) with 0.13 mm sensor diameter was inserted through the renal parenchyma into the calyceal lumen, adjacent to the stone tissue boundary for temperature monitoring. The distance between the thermocouple tip and the fiber/stone interface was standardized as much as possible across experiments.

Each renal unit was each assigned to saline or 0.5 wt.% ITO@SiO_2_ in saline for irrigation. Continuous room-temperature irrigation was run at a flow rate of 20 mL min^-1^ using a peristaltic pump (Maserflex®). LL was performed using a Ho:YAG system using a 270-µm fiber. In the kidney, LL employed a dusting technique aiming to reduce kidney stone to particles smaller than 0.25 mm. LL was performed at 0.4 J and 20 Hz using a 50% duty cycle (i.e., 30 s on/30 s off), with total lasing duration of 3 min. The laser was stripped between duty cycles (every 1 min). Sham procedures replicated the full intervention without laser activation and consisted of ureteroscopy with temperature monitoring adjacent to the stone for a duration of 5 min. Endoscopic video footage was recorded. Upon completion of the bilateral procedures, the pigs were euthanized, and bilateral nephrectomies were performed for histopathological examination.

#### Histopathological analysis

The kidneys and ureters were grossly inspected for evidence of thermal injury and necrosis. Tissue sections from each specimen were taken to include cross sections of the lumens of renal pelvis/calyces, ureters, and renal parenchyma. These were formalin-fixed and embedded in paraffin. Slides were cut from the paraffin tissue blocks and stained with hematoxylin and eosin. The slides were examined microscopically by an experienced genitourinary pathologist (J.H.) for the presence or absence of thermal injury. Mechanical injury was characterized by denuded urothelium, focal hemorrhage, and vascular congestion. Minor thermal injury was defined as focal or superficial collagen homogenization (<1 mm), while major thermal injury was defined as by global and deep collagen homogenization (>1 mm), fibrin deposition, edema, and loss of cellular details.

#### Inductively coupled plasma mass spectrometry (ICP-MS)

All samples were diluted using a 3 wt.% aqueous HNO_3_ matrix. Calibration curves were generated via serial dilutions of a 10 ppm In^3+^ multi-element standard (Ricca Chemical, 5% HNO_3_). To prepare the control sample, 0.5 wt.% ITO@SiO_2_ was digested in 1 M HNO_3_ at 60 °C for 2 days; the resulting supernatant was centrifuged and diluted 10^4^ to 10^6^-fold to yield element concentrations between 1 ppb and 1 ppm. The untreated sample was prepared by directly diluting the supernatant of the centrifuged 0.5 wt.% ITO@SiO_2_ dispersion. For the laser-treated group, LL (*E_p_* = 0.4 J, *f* = 20 Hz) was applied to 20 mL of the 0.5 wt.% dispersion for 1 min, followed by the identical centrifugation and dilution procedure. Analyses were performed on either a Thermo iCAP Q or Thermo iCAP RQ ICP-MS.

#### Cytotoxicity test

Cell culture: A murine epithelial cell line (mIMCD-3, ATCC, Manassas, VA, USA) was used for all experiments. Cells were cultured in Dulbecco’s Modified Eagle Medium/Hams F-12 (DMEM/F-12; pH 7.4, #11 320 033, Thermo Fisher, Waltham, MA, USA), supplemented with 10% fetal bovine serum (FBS, #10 437 028, Thermo Fisher, Waltham, MA, USA). Cells grew at 5% CO_2_ at 37 °C and passaged upon reaching 80% confluency with a maximum passage number of 20.

MTT Assay: MIMCD-3 cells were seeded onto sterile 12-well plates (#3513, Corning Inc, Corning NY, USA) at a density of 100,000 cells per well. The cells were cultured overnight and incubated with untreated and treated ITO@SiO_2_ (0.5 wt.%) for 1 h or 24 h. Control cells were incubated in DMEM/F-12 media for the duration of the treatment. Incubation with Triton X-100 (10% in Dulbecco’s Phosphate Buffered Saline (PBS), 30 s exposure; X100-100ML, Millipore Sigma, Burlington, MA, USA) was used to damage cells, serving as positive control. Cells were washed 3 times in PBS (#28 374, Thermo Fisher, Waltham, MA, USA). Cells were incubated in DMEM (300 µl; #31 053 028, Thermo Fisher) and 3-(4,5-dimethylthiazol-2-yl)-2,5-diphenyl tetrazolium bromide (MTT, 0.5 mg mL-1, #V13154, Thermo Fisher, Waltham, MA, USA) for 1 h. Media was aspirated, and cells were incubated in dimethyl sulfoxide (300 µl; DMSO, #D8418, Millipore Sigma) for 10 min in the dark. DMSO was transferred into a clear 96-well plate (#82 050, VWR, Radnor, PA, USA) and absorbance was measured at 580 nm using a plate reader (SpectraMax iD3, Molecular Devices, San Jose, CA, USA).

## Supporting Information

Supporting Information is available from the author.

## Acknowledgements

The authors acknowledge the financial support from NIH National Institute of Diabetes and Digestive and Kidney Diseases (5P20-DK135107-03 and 5R01DK138972-02). Q.F. acknowledges the helpful discussion with Jinxing Chen, Yichen Li and Yadong Yin in the nanoparticle synthesis.

## Author Contributions

Q.F. and J.C. contribute equally to this work. P.-C.H. and P.Z. conceived the study and was responsible for project management. Q.F. performed the nanoparticle synthesis and optical measurements on nanofluid. J.C. performed the stone damage assessment and collected bubble dynamics data. A.M. coordinated the hydrogel kidney and porcine laser lithotripsy studies, contributed to experimental planning and execution, and assisted in data collection, and A.S. performed the LL in the hydrogel kidney model. M.B. performed the LL in the porcine kidney. J.D. and C.C. performed the cytotoxicity experiment. J.L. performed the ICP-MS test. Y.C., R.W., T.-H.C., C.P., and M.L. were involved in conceptualization of the study. Q.F. analyzed and visualized all the data and drafted the manuscript with P.-C.H. All other authors provided critical feedback and helped shape the study and edit the manuscript.

---

**Extended Data Fig. 1.**
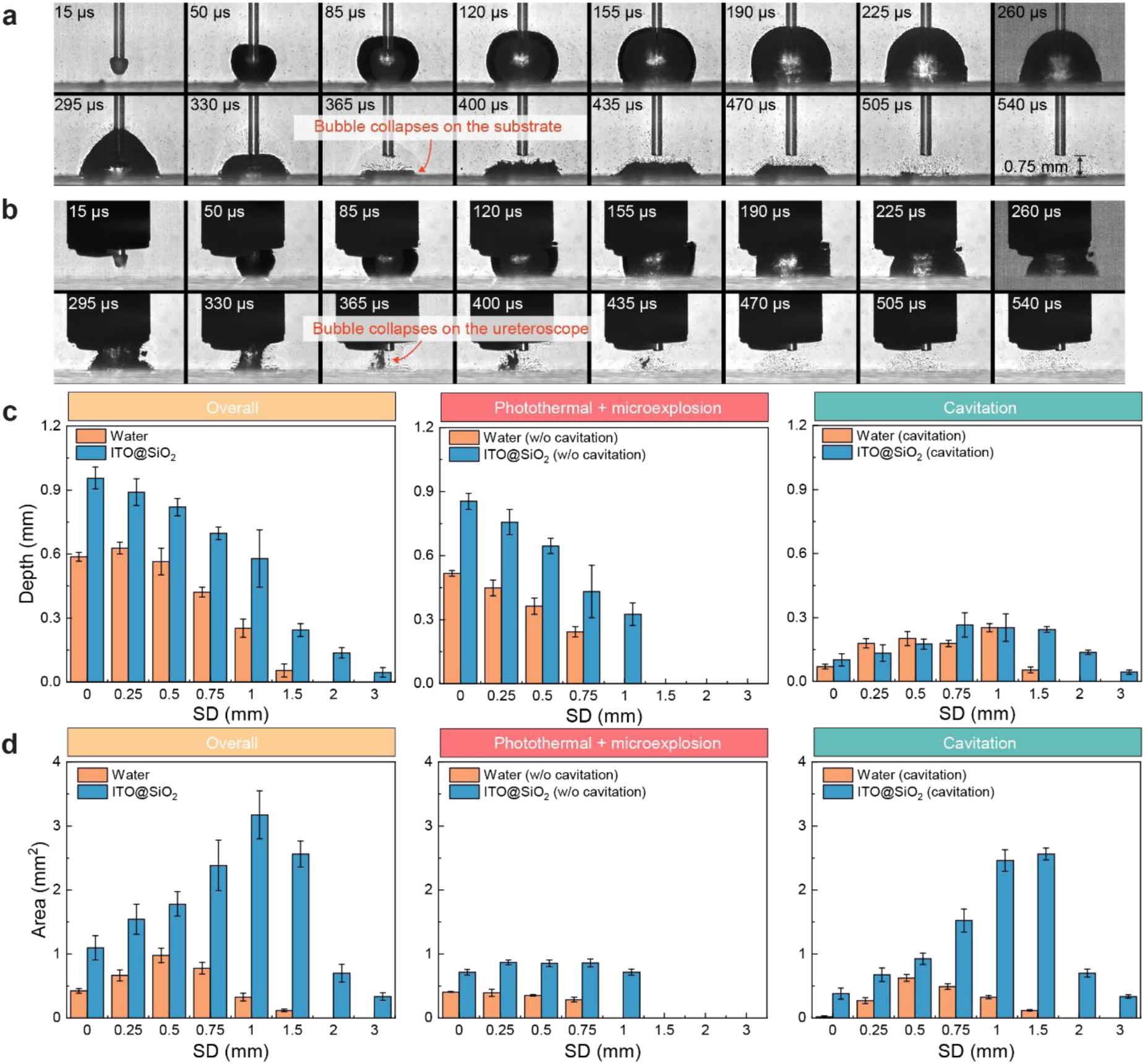
(a, b) High-speed digital photographs demonstrating the shift in the bubble collapse site—from the substrate (a) to the ureteroscope (b)—achieved by reducing the distance between the ureteroscope and the fiber tip from 5 mm to 0.25 mm. (c, d) Decoupled contributions of photothermal/microexplosion effects versus cavitation damage to the resulting crater depth (c) and profile area (d) across various standoff distances (SDs).

**Extended Data Fig. 2.**
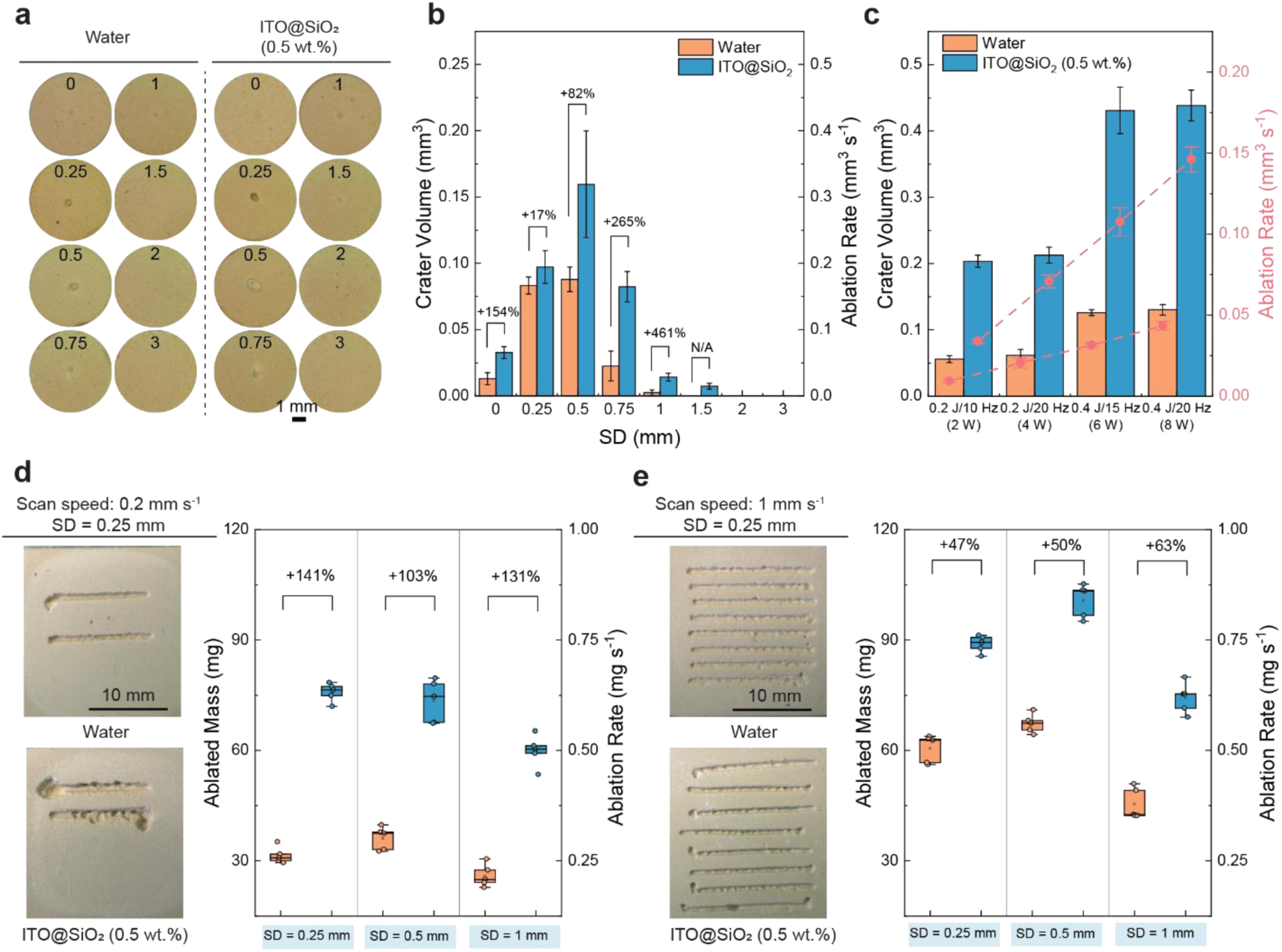
(a) Digital photographs of craters generated via spot treatment (*E_p_* = 0.4 J, *f* = 20 Hz, *PN* = 10) in water and 0.5 wt.% ITO@SiO_2_. Inset numbers denote the specific SD values. (b) Corresponding crater volumes from panel (a). (c) Crater volumes and ablation rates achieved at various power settings during spot treatment in water and 0.5 wt.% ITO@SiO_2_ (SD = 0 mm, *PN* = 60). (d, e) Scanning treatment results at scan speeds of 0.2 mm s^-1^ (d) and 1 mm s^-1^ (e) with a 2-min lasing duration. For both panels, the left images display representative photographs of the BegoStone slabs following scanning treatment at an SD of 0.25 mm. (Note: Optical fibers were cleaved every minute to minimize the impact of fiber degradation under both conditions).

**Extended Data Fig. 3.**
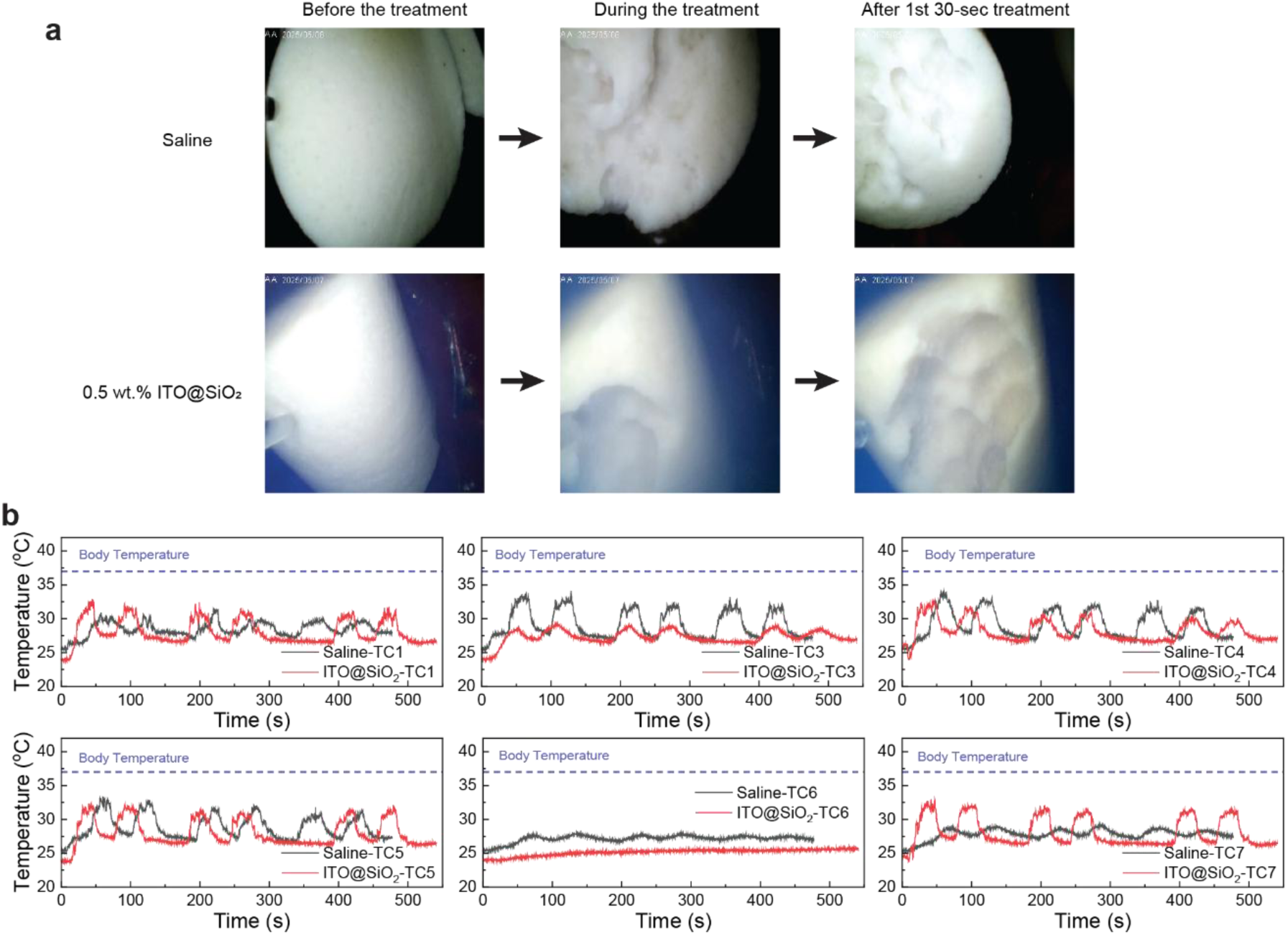
(a) Ureteroscopic views of BegoStones before, during, and after the initial 30-second treatment period in saline and the 0.5 wt.% ITO@SiO_2_ nanofluid. (b) Temperature profiles recorded by the remaining thermocouples during the hydrogel kidney model experiment.

**Extended Data Fig. 4.**
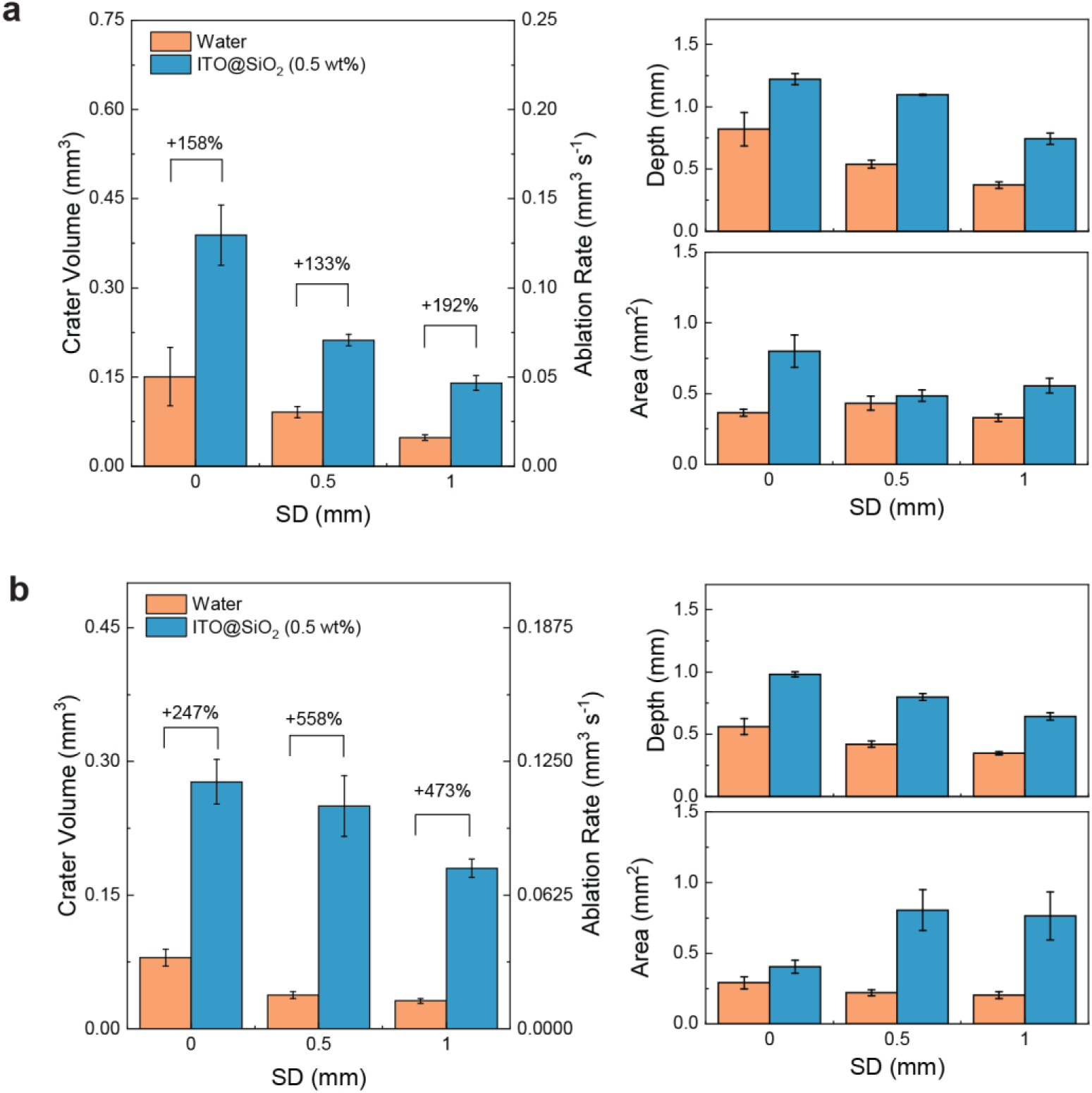
Ablation efficiency assessment in a bench-top cuvette model for TFL and Tm:YAG lasers. (a) Volume, depth, and profile area of craters generated in water and 0.5 wt.% ITO@SiO_2_ via TFL spot treatment (*E_p_* = 0.2 J, *f* = 20 Hz, *PN* = 60). (b) Volume, depth, and profile area of craters generated in water and 0.5 wt.% ITO@SiO2 via Tm:YAG spot treatment (*E_p_* = 0.2 J, *f* = 25 Hz, *PN* = 60).

