## Supplementary information for "Visibly Transparent, Near-Infrared Absorbing Nanofluids Enable High-Efficiency and Safe Laser Lithotripsy"

| | STONE TRACKING | RECURRENCE RATE[1] | COMPLICATION RATE[2] | SICK LEAVE DAYS[1] | HOSPITAL COSTS[1*] | CONTRAINDICATIONS[2] |
|---|---|---|---|---|---|---|
| **ESWL** | X-ray/ultrasound (indirect vision) | 26.2% | 1-4% (ureter) | 10.1 | €3240 | Pregnancy, bleeding diatheses, anatomic abnormalities, cystine and uric acid stones |
| **URS** | Endoscope (direct vision) | 15.3% | 7-11% (ureter) | 6.8 | €2979 | Active/untreated infection |
| **PCNL** | Endoscope (direct vision) | 22.9% | 15% (kidney) | 13.0 | €5783 | Pregnancy, bleeding diatheses, active infection, Anatomic constraints |

**Supplementary Table 1.** Comparison of three modalities of urinary stone treatment (*total hospital cost including re-interventions).

| STONE TYPE/LOCATION | ESWL | URS | PCNL |
|---|---|---|---|
| Ureteral stones (overall) | 72% | 90% | - |
| Renal pelvic stones (>20 mm) | - | 75% | 94% |
| Lower pole renal stones (10-20 mm) | 58% | 81% | 87% |
| Lower pole renal stones (>20 mm) | 10% | 83% | 71% |
| Staghorn stones | 19-57% | - | 69.5-76.9% |

**Supplementary Table 2.** Comparison of Stone-Free Rates for SWL, URS, and PCNL. Data from Ref [2].

| LITHOTRIPTER | TYPE | YEAR OF INVENTION | ADVANTAGES | DISADVANTAGES |
|---|---|---|---|---|
| **Pneumatic**[3–6] | Compressed air | 1800s | Cost effective | Strong retropulsion<br>Risk of perforation<br>Mostly rigid probe |
| **Ultrasonic**[3,5–7] | High-frequency sound waves | 1950s | Cost effective<br>Durable probe<br>Simultaneous evacuation of fragments | Rigid, large probe<br>Strong retropulsion |
| **Electrohydraulic**[3,5–7] | Shock wave generated by electric current | 1950s | Cost effective<br>Flexible, small probe<br>Effective at breaking all stones | Narrow margin of safety<br>Rapid degradation of probe<br>Strong retropulsion |
| **Pulsed ruby laser**[8,9] | Laser (694 nm) | 1968 | Solid-state laser | High risk of thermal injury<br>*Never in clinical use |
| **Q-switched Nd:YAG laser**[3,7,9] | Laser (1064 nm) | 1983 | Solid-state laser | Inability to break calcium oxalate monohydrate and brushite stones<br>Fragile delivery system<br>Large fiber (600 µm)<br>Low pulse energy |
| **Pulsed dye laser**[3,6,7] | Laser (504 nm) | 1986 | Wide margin of safety<br>Small fiber (200-400 µm) | Inability to break calcium monohydrate and cystine stones<br>Liquid laser |
| **Alexandrite laser**[6,10] | Laser (750 nm) | 1988 | Solid-state laser | High cost<br>Inability to break calcium monohydrate and cystine stones |
| **Ho:YAG laser**[3–5,9] | Laser (2120 nm) | 1993 | Solid-state laser<br>High pulse energy<br>Small fiber (200-1000 µm)<br>Effective at breaking all stones | High cost<br>Fast fiber degradation at high energy settings<br>Risk of thermal injury<br>High cost |
| **Erbium:YAG laser**[11] | Laser (2940 nm) | 2001 | Solid-state laser<br>Higher absorption | Lack of a suitable fiber delivery system |
| **Thulium fiber laser**[11] | Laser (1940 nm) | 2005 | Solid-state laser<br>Higher absorption<br>Air cooling | Lack of sufficient clinical data |
| **Tm:YAG laser**[11] | Laser (2010 nm) | 2015 | Solid-state laser<br>Higher absorption | Lack of sufficient clinical data |

**Supplementary Table 3.** The comparison of various Intracorporeal lithotripters in URS. (Solid-state laser has the advantage of better reliability and easier maintenance compared to the liquid laser).

| REF. | MATERIAL ADDED IN THE FLUID | LASER WAVEL ENGTH | BUBBLE DYNAMIC S | STONE DAMAGE ASSESSMENT | TOXICITY | ADDITIONAL NOTE |
|---|---|---|---|---|---|---|
| [12] AND [13] | Polyhydroxy fullerene/ multiwalled carbon nanotubes/Au nanospheres/A u nanorods/ graphene oxide | 785/1320 nm (Continu ous wave) | Not investigated | Benchtop model | Not investigated | Not clinically relevant (LL was done in air), Pretreatment of stone |
| [14] | Potassium dichromate | 504 nm | With stone | Benchtop model | Highly toxic | |
| [15] AND [16] | Tween 80/Tween 20 | 1940 nm (Thulium fiber laser) | Without stone | Not investigated | Not investigated | Changing the shape of bubbles by surfactants |
| [17] | Hollow Prussian bule nanoparticles | 808 nm (Continu ous wave) | Not investigated | Ex vivo | Good biocompatib ility | Not clinically relevant (LL was done in air), Pretreatment of stone |
| [18] | PEDOT:PSS | 2120 nm (Ho:YA G) | With stone | Benchtop model | Good biocompatib ility for moderate concentratio ns | Use of clinical Ho:YAG laser, Compatible with clinical practice |
| THIS WORK | ITO@$SiO_2$ nanoparticles | 2120 nm (Ho:YA G) | With stone | In vivo | Good biocompatib ility | Use of clinical Ho:YAG laser, Compatible with clinical practice |

**Supplementary Table 4.** Comparison of various approaches that utilize nanoparticles/dyes/surfactants in laser lithotripsy. (Continuous wave lasers created too much heat and caused tissue damage and were inappropriate for use for stone fragmentation.[19])

**Supplementary Note 1. Morphologies of ITO nanoparticles of various feeding ratios**

The size and shape of the ITO nanoparticles remain largely unchanged as the feeding ratio increases from 0 to 6.25%. However, as the feeding ratio further increases to 7.69%, the shape of the nanoparticles becomes less cuboidal. In addition, the absorption peak intensity shows a slight decrease. This is presumable due to the interruption of the Sn atoms to the crystal growth of $In_2O_3$ at high feeding levels.

| Feeding ratio | 0% | 3.23% | 6.25% | 7.69% |
|---|---|---|---|---|
| Avg. part. size | 18.2 ± 3.8 nm | 16.9 ± 3.4 nm | 19.6 ± 4.1 nm | 18.1 ± 4.6 nm |

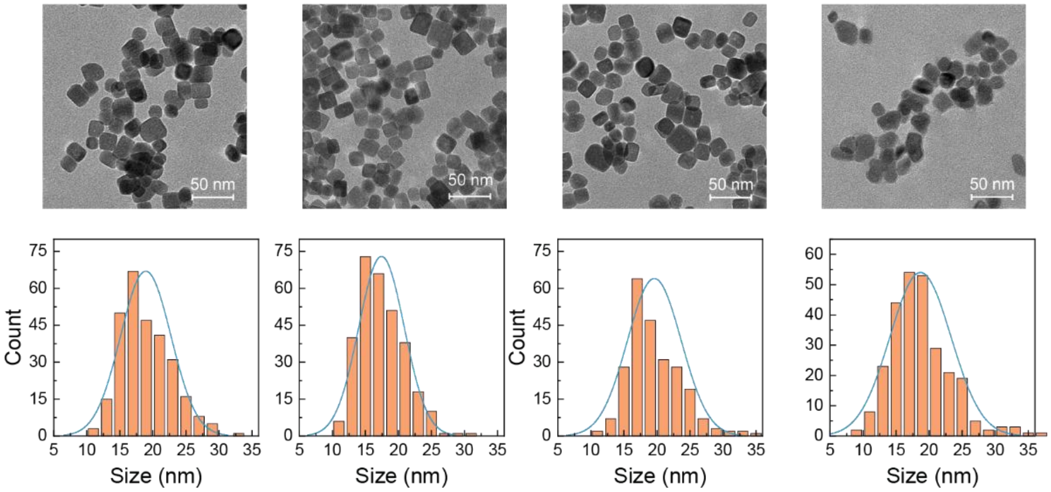

**Supplementary Fig. 1.** The morphologies and size distributions of ITO nanoparticles synthesized with different doping ratios.

**Supplementary Note 2. Colloidal stability of nanoparticles before and after surface modification**

The hydrodynamic diameters of the nanoparticles were measured on Malvern Zetasizer Nano ZS five times and their average values and standard deviations were plotted in the figure below. All the nanoparticles have hydrodynamic diameters larger than their size in TEM due to the presence of the solvation layer.

Although the grafting of PA on ITO nanoparticles increases their colloidal stability in saline or PBS, the solutions quickly become blurry upon the addition of BegoStones. Conversely, the ITO@$SiO_2$ solutions stay clear for days even with the presence of BegoStones. Therefore, the colloidal stability of the nanoparticles is: ITO@$SiO_2$ > ITO-PA > ITO.

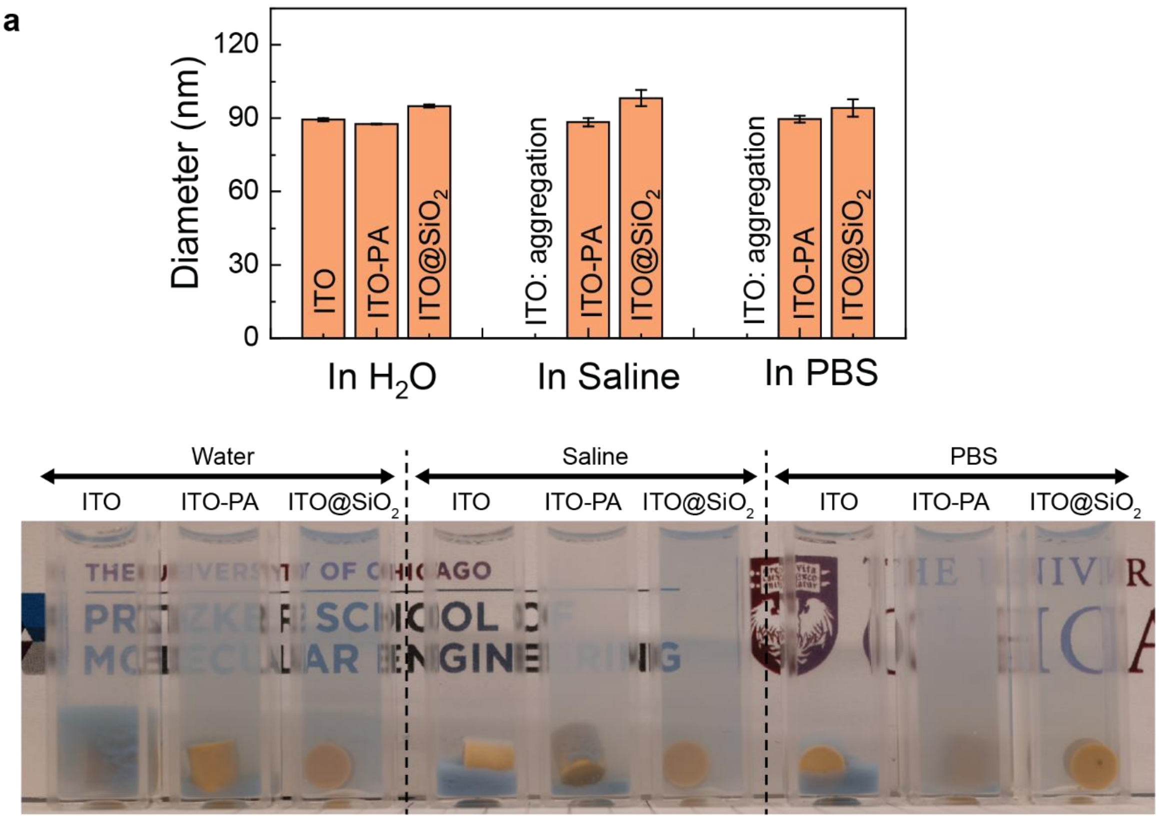

**Supplementary Fig. 2.** (a) Hydrodynamic diameters of the nanoparticles before and after surface modification in water, saline and PBS. (b) Digital photographs demonstrating colloidal stability of the nanoparticles before and after surface modification in the presence of BegoStones.

**Supplementary Note 3. Optical properties of ITO and ITO@$SiO_2$ nanoparticles**

After the $SiO_2$ coating, the absorption intensity decreases slightly due to the reduced effective concentration of ITO. Meanwhile, the peaks show a 50-100 nm redshift compared to the bare ITO nanoparticles. This is due to the higher refractive index of the $SiO_2$ shell that shifts the localized surface plasmon resonance of ITO to longer wavelengths.

For ITO@$SiO_2$ with the feeding ratio of 4.76%, at the concentration of 0.5 wt.%, its absorption coefficient is 0.0126 $\mu m^{-1}$ (penetration depth: 79 μm), corresponding to a 394% increase compared to that of water (0.00255 $\mu m^{-1}$, penetration depth: 392 μm).

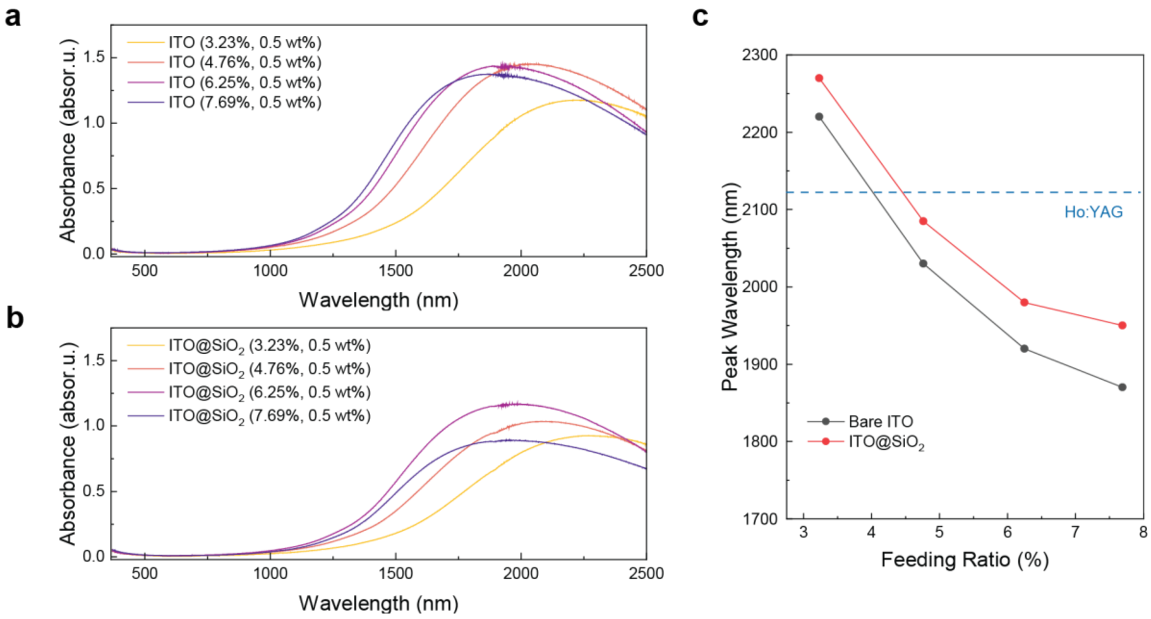

**Supplementary Fig. 3.** (a, b) Vis-NIR absorption spectra of bare ITO (a) and ITO@$SiO_2$ (b) at various feeding ratios (water baseline). (c) The relationship between the peak wavelength and feeding ratio for bare ITO and ITO@$SiO_2$ nanoparticles.

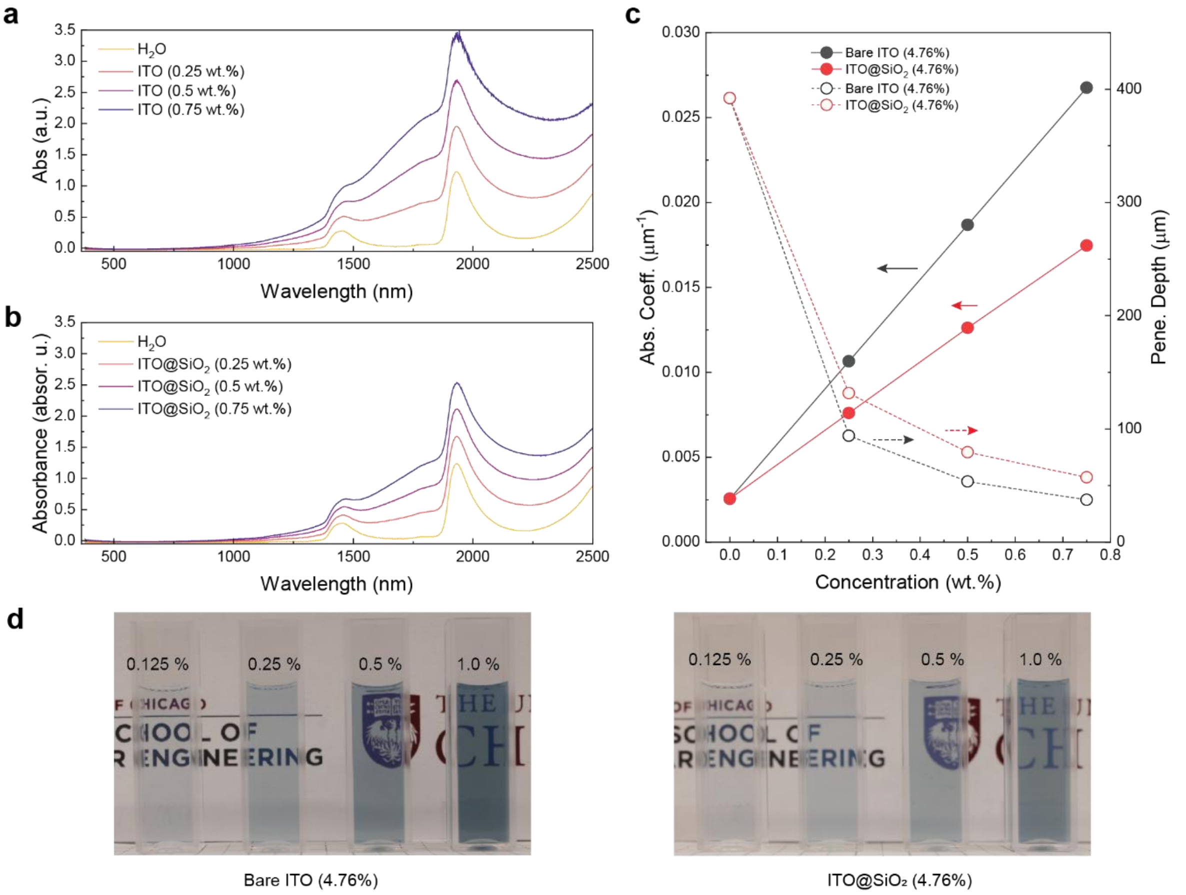

**Supplementary Fig. 4.** (a, b) Vis-NIR absorption spectra of bare ITO (a) and ITO@$SiO_2$ (b) of various concentrations (air baseline, feeding ratio: 4.76%). (c) Absorption coefficient and penetration depth of bare ITO and ITO@$SiO_2$ at different concentrations (feeding ratio: 4.76%). (d) Digital photographs of the ITO and ITO@$SiO_2$ nanoparticle solutions of various concentrations (path length: 1 cm).

**Supplementary Note 4. Synthesis of $SiO_2$, carbon, and gold Nanoparticles**

$SiO_2$ nanospheres with a diameter of ~75 nm were synthesized using the classical stober method.[20] 196 mL ethanol, 20 mL Millipore water, and 2 mL $NH_4OH$ (28 wt %) were mixed in a 250 mL flask. After the mixture was magnetically stirred for 5 min, 10 mL of TEOS was injected. The mixture was maintained at room temperature for 15 h with stirring (300 rpm). The nanospheres were separated from the reaction mixture by centrifuging at 11000 rpm for 15 min. The product was further purified by dispersing the nanospheres in ethanol with the help of sonication bath and centrifuging (11000 rpm for 15 min). The same purification process was repeated using water as the solvent for twice more. The final product of $SiO_2$ nanoparticles was dispersed in water and its concentration was determined by drying a certain amount of solutions in the oven.

Carbon nanoparticles were synthesized by etching the carbon in NaOH solution. 0.2 g NaOH was dissolved in 50 mL water to make 0.1 M NaOH solution. 1 g of biocarbon was added to the above NaOH solution and the mixture was heated to 70 °C on a hotplate with constant magnetic stirring for 3 days. After cooling to the room temperature, the mixture was centrifuged using 8500 rpm for 15 min and the solid was discarded. The supernatant was titrated with 1 M HCl solution to adjust its pH into ~7. The brownish solution was transferred to a semi-permeable membrane (cutoff 14 kDa) and dialyzed in water for 2 days to remove the NaCl. The concentration of C nanoparticle solution was determined by drying a certain amount of solution in the oven. Because the C nanoparticles were prepared by an uncontrolled etching method, their morphology is not well defined.

Au nanoparticles with a diameter of ~30 nm were synthesized using a two-step seeded growth method.[21] Au seeds were prepared first by directly reducing the $HAuCl_4$ using sodium borohydride. 1 mL of $HAuCl_4$ (5 mM) and 1 mL of TSC (5 mM) were mixed with 18 mL of $H_2O$ in a flask. Under vigorous stirring, 0.6 mL of freshly made $NaBH_4$ solution (0.1 M) was quickly injected into the solution, leading to an immediate color change to yellowish red. After stirring for 4 h, the solution was collected as the seed solution for subsequent seeded growth. The second step involves the growth of Au on the premade Au seeds. under vigorous stirring, 200 μL of the seed solution was then quickly injected into a freshly prepared growth solution of Au containing 500 μL of PVP (5 wt.%), 250 μL of l-ascorbic acid (0.1 M), 200 μL of KI (0.2 M), 60 μL of $HAuCl_4$ (0.25 M), and 2 mL of $H_2O$. After 10 min the Au nanoparticles formed were collected by centrifugation and redispersed in water. PVP, the surface ligand, can effectively increase the colloidal stability of Au nanoparticles against aggregation.

It is difficult to image the PEDOT:PSS nanoparticles under transmission electron microscope, because of the dehydration and deformation of PSS segments during drying.

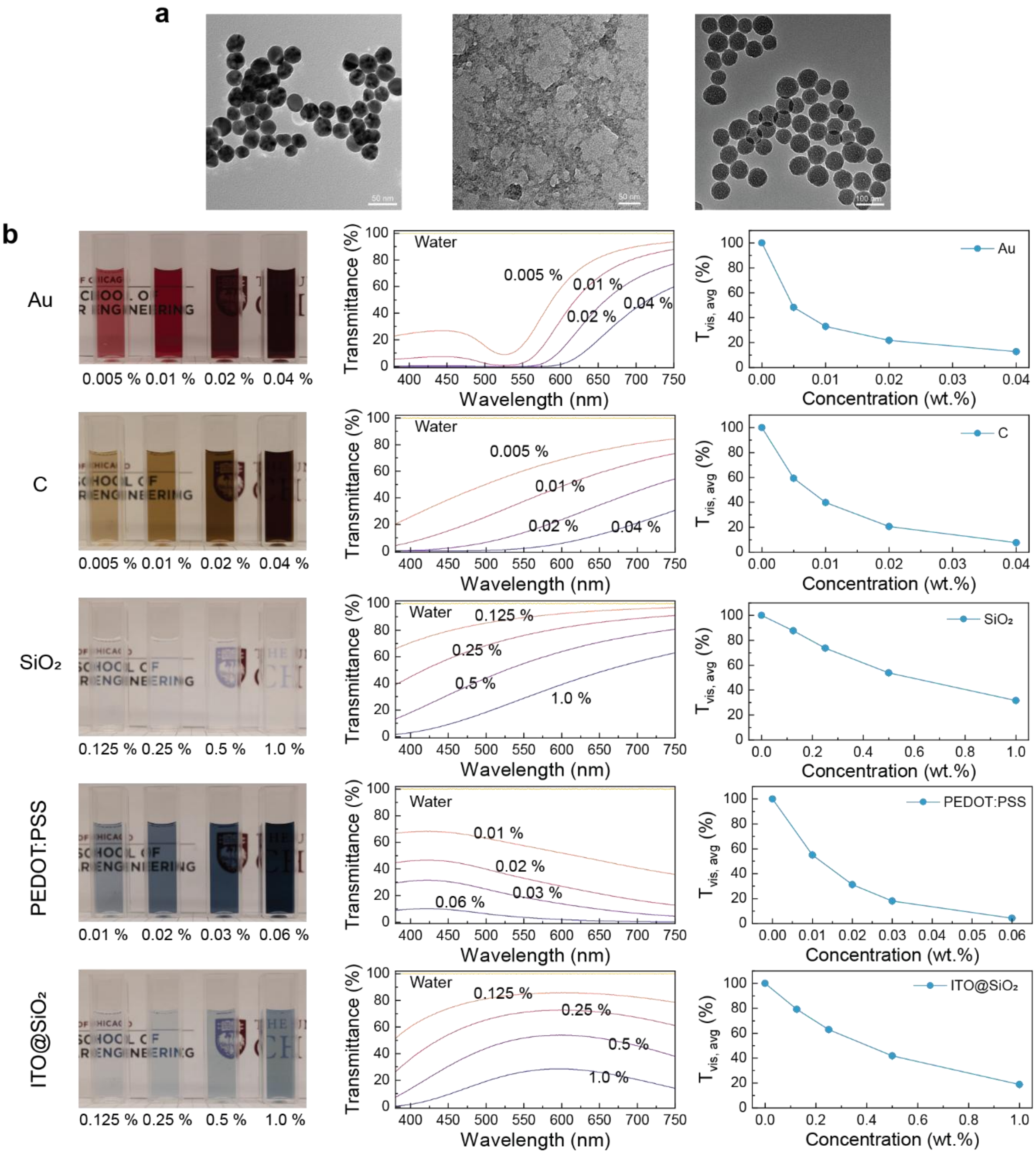

**Supplementary Fig. 5.** (a) TEM images of Au, C and $SiO_2$ nanoparticles. (b) Digital photos of various nanoparticle solutions, measured visible transmittance, and calculated average visible transmittance of nanoparticles solutions with various concentrations.

**Supplementary Note 5. Ureteroscopic views of BegoStones in nanoparticle solutions**

When taking the photo using ureteroscope, BegoStones were immersed in fluids in the cuvette, and the whole cuvette was wrapped with a black foil to block the ambient light. The offset distance between the fiber tip and the ureteroscope was set to 5 mm in the visibility test and there was an additional 2 mm set for the SD, so the distance between the ureteroscope and the stone surface in the images below was 7 mm in total.

To quantitively analyze the visibility of ureteroscope in various nanofluids, we selected an area of 1000*1000 $pixel^2$ (red box) on the stone and three areas of 500*500 $pixel^2$ (green boxes) in the surrounding (**Supplementary Fig. 7**) and calculated the relative luminance (defined by Web Content Accessibility Guidelines 2) of each pixel based on the following equation:

$$relative\ luminace = 0.2126 * R + 0.7152 * G + 0.0722 * B$$

where R, G, and B are defined as:

- if RsRGB <= 0.03928 then R = RsRGB/12.92 else R = ((RsRGB+0.055)/1.055) ^ 2.4
- if GsRGB <= 0.03928 then G = GsRGB/12.92 else G = ((GsRGB+0.055)/1.055) ^ 2.4
- if BsRGB <= 0.03928 then B = BsRGB/12.92 else B = ((BsRGB+0.055)/1.055) ^ 2.4

and RsRGB, GsRGB, and BsRGB are defined as:

RsRGB = R8bit/255; GsRGB = G8bit/255; BsRGB = B8bit/255.

The difference between the stone area and the surrounding areas is defined as the relative luminance contrast.

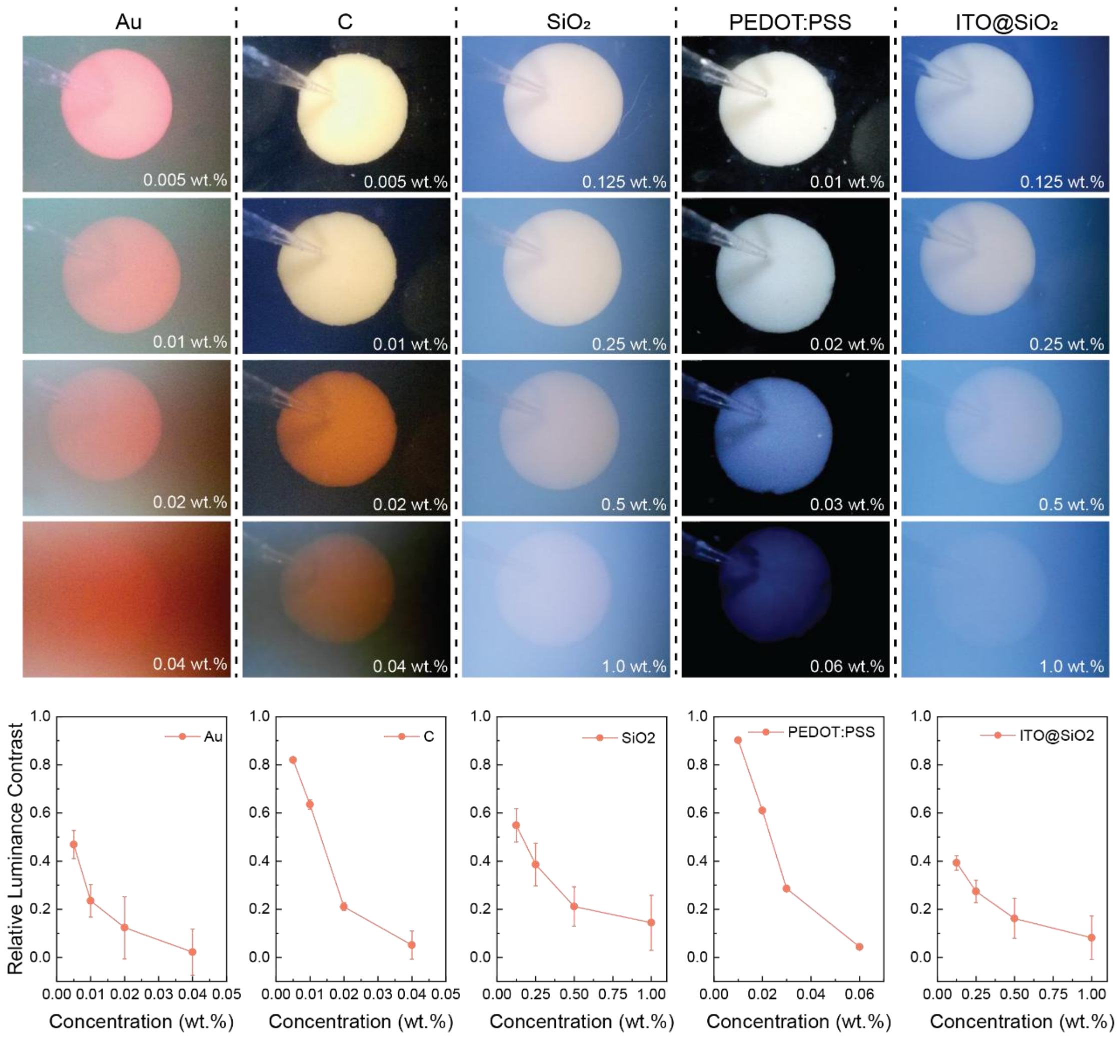

**Supplementary Fig. 6.** Ureteroscopic views of BegoStones immersed in the nanoparticle solutions with different concentrations.

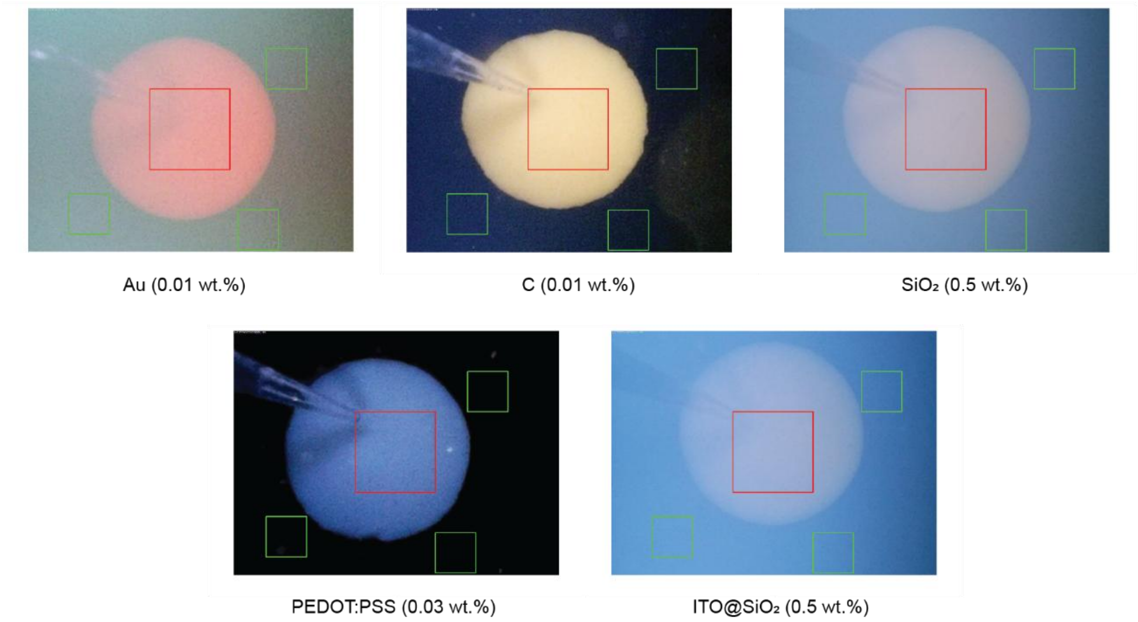

**Supplementary Fig. 7.** Images showing the selection of stone area (red box) and surrounding areas (green boxes) for the calculation of relative luminance**.**

**Supplementary Note 6. The calculation of transmitted power**

Time-dependent transmittance is calculated based on the Beer-Lambert law by considering the measured temporal distance:

$$T(t) = \frac{I(t)}{I_0} = e^{-\alpha_v l_v(t) - \alpha_l l_l(t)}$$

where $\alpha_v$ and $\alpha_l$ are the absorption coefficient of the vapor and liquid phase. $l_v$ and $l_l$ are the light path in the vapor and liquid phase from the fiber tip to the detector at a specific standoff distance. The sum of $l_v$ and $l_l$ was equal to the SD at each setting. In the calculation, the $\alpha_v$ is set to be 0.000001μm$^{-1}$. And the $\alpha_l$ values are 0.00255 μm$^{-1}$ and 0.0126 μm$^{-1}$ for water and 0.5 wt.% ITO@$SiO_2$ respectively.

The transmitted power is calculated by multiplying the laser output power profile with the time-dependent transmittance. And the transmitted energy (area enclosed by the transmitted power curve) is obtained by integrating the transmitted power with respect to the time. At the SD ≤ 0.75 mm, the transmitted energy in 0.5 wt.% ITO@$SiO_2$ is smaller than that in water. This is because although the bubbles expand faster in ITO@$SiO_2$, the short penetration depth leads to a higher energy loss at the beginning of the laser activation. But as the SD further increases (≥ 1 mm), the additional energy transmitted due to the faster bubble expansion starts to exceed the initial energy loss due to the short penetration depth. And the larger the SD, the larger the difference.

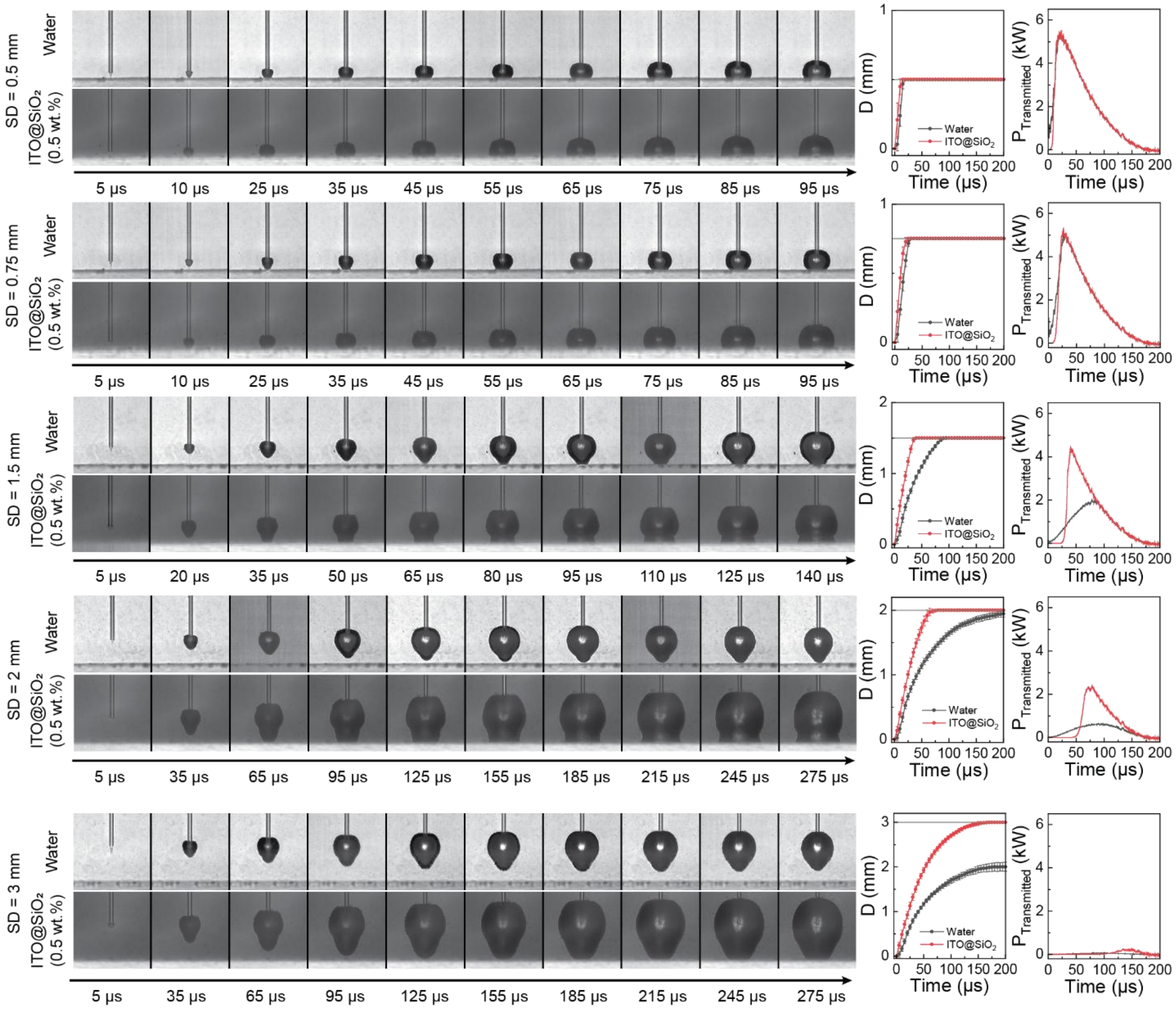

**Supplementary Fig. 8.** High-speed imaging of bubble expansion in front of a glass substrate at various SDs in water and the 0.5 wt.% ITO@SiO$_2$ solution ($E_p$ = 0.4 J). And measured apex-to-fiber distance, calculated laser power transmitted through the glass substrate at the corresponding SD.

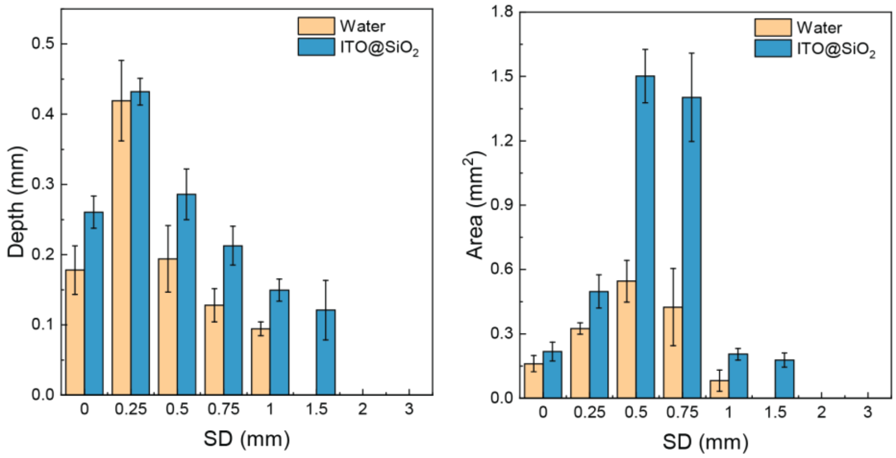

**Supplementary Fig. 9.** Depth, and profile area of the craters generated in water and 0.5 wt.% ITO@$SiO_2$ solutions in the spot treatment ($E_p$ = 0.4 J, $f$ = 20 Hz, $PN$ = 10).

**Supplementary Note 7. Additional H&E-stained histological sections of the renal tissues**

Similar to the H&E-stained sections of renal tissues in the main text, both sham groups show normal histological morphology without the sign of thermal or mechanical injury. However, in the treated groups, except the mechanical injury such as the denudation of the urothelium, additional hemorrhage is observed in Pig 2, although it is regional and relative mild.

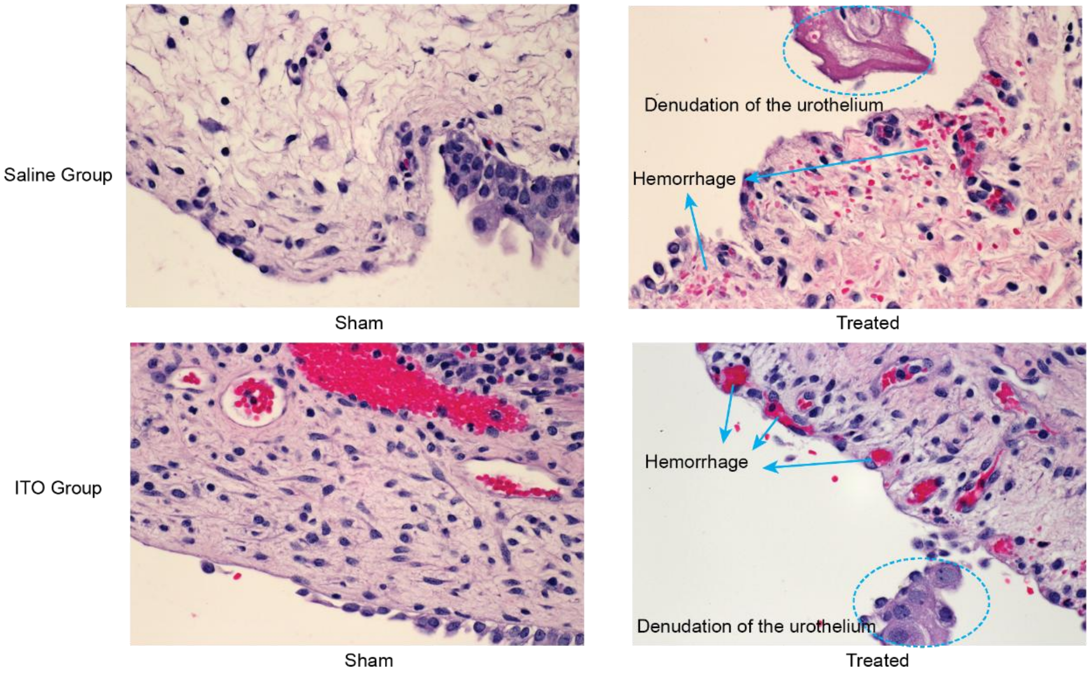

**Supplementary Fig. 10.** Representative hematoxylin and eosin (H&E)-stained histological sections of the renal tissues of Pig 2.

**Supplementary Note 8. ICP-MS test**

Eight standard $In^{3+}$ solutions were made by diluting the stock solution containing 10 ppm (10 µg $mL^{-1}$) $In^{3+}$ using 3% $HNO_3$ solution into 1000, 500, 200, 100, 50, 10, 5, and 1 ppm. The sample dilutions were made with a step of 10 times, and those with concentration within 1-1000 ppb are included in the table.

| Sample | Dilution | [$In^{3+}$] measured by ICP (ppb) | [$in^{3+}$] in original solution (ppb) | [$In^{3+}$] (µg $mL^{-1}$) |
|---|---|---|---|---|
| Control | $10^4$ | 278.95209769644 | 2789520.98 | 2815.80 ± 18.77 |
| | $10^5$ | 28.2566457003643 | 2825664.57 | |
| | $10^6$ | 2.83220545225904 | 2832205.45 | |
| Untreated | 10 | 196.0376128135 | 1960.38 | 1.99 ± 0.08 |
| | $10^2$ | 19.1684880507881 | 1916.85 | |
| | $10^3$ | 2.10185887009792 | 2101.86 | |
| Treated | 10 | 247.960444545513 | 2479.60 | 2.42 ± 0.05 |
| | $10^2$ | 23.4713162718878 | 2347.13 | |
| | $10^3$ | 2.43004235520814 | 2430.04 | |

**Supplementary table 5.** Concentrations of $In^{3+}$ in supernatants with various dilutions.
